# Aromatic Ring Flips Reveal Reshaping of Protein Dynamics in Crystals and Complexes

**DOI:** 10.1101/2025.08.20.671406

**Authors:** Lea M. Becker, Haohao Fu, Ben P. Tatman, Matthias Dreydoppel, Anna Kapitonova, Ulrich Weininger, Sylvain Engilberge, Christophe Chipot, Paul Schanda

## Abstract

Protein conformational energy landscapes are shaped not only by intramolecular interactions but also by their environment. In protein crystals and protein-protein complexes, intermolecular contacts alter this energy landscape, but the exact nature of this alteration is difficult to decipher. Understanding how the crystal lattice affects protein dynamics is crucial for crystallography-based studies of motion, yet its influence on collective motions remains unclear. Aromatic ring flips in the hydrophobic core represent sensitive probes of such dynamics. Here, we compare the kinetics of aromatic ring flips in the protein GB1 in crystals, in complex with its binding partner IgG, and in solution, combining advanced isotope labeling with quantitative NMR methods. We show that rings in the core flip nearly a thousand times less frequently in crystals than in solution. Enhanced-sampling molecular dynamics simulations, based on a new crystal structure, reproduce these elevated barriers and reveal how the crystal restrains motions.

## Introduction

Protein crystallography has been the central pillar of structural biology for over half a century, with two hundred thousand structures determined up to now.^1^ Notwithstanding the enormous insight they provide, it has been recognized early that the seemingly static images provided by crystallography – obtained primarily at cryogenic temperatures – are misleading; as G. Weber expressed already in 1975, proteins are “kicking and screaming stochastic molecules” in solution.^2^ While the seemingly rigid crystal structures could lead to the assumption that the crystal quenches dynamics, it has become clear that proteins in crystal lattices still exhibit significant motion. For example, it is common to find motion-related residue-by-residue variability of B-factors or even complete blurring of the electron density. Distinct densities arising from co-existing conformations^3,4^ (so-called alternate locations) are found in thousands of crystal structures.^5^ As data collection at room temperature is becoming more widely used thanks to serial crystallography,^6–8^ the observation of local dynamics in crystals is becoming increasingly common. ^9,10^

A long-standing question, posed more than half a century ago, is to what extent crystal packing constrains protein dynamics relative to solution.^11,12^ Early studies of enzymatic activity in crystals^13–19^ revealed contrasting outcomes: in some cases, such as oxygen binding in hemoglobin, crystal properties closely matched those in solution,^20^ whereas in others, catalytic activity was strongly attenuated.^13^ Structural comparisons of proteins in different crystal forms further established that packing can alter both structure and dynamics.^21–23^ This issue has gained renewed urgency as serial crystallography at X-ray free-electron lasers and synchrotrons allows capturing protein kinetics at a time resolution down to femtoseconds, thereby allowing to probe protein mechanisms and reaction intermediate states of protein crystals.^7,8,24–30^ These experiments inherently build upon the assumption that the crystal lattice does not influence protein dynamics.

Directly assessing how the lattice modifies protein dynamics remains challenging. Crystallographic indicators of heterogeneity, such as B-factors, conflate static disorder and thermal motion; they do not provide information on timescales, and naturally cannot assess dynamics of non-crystalline samples. Consequently, crystallography alone cannot quantify the extent to which the lattice constrains protein motions relative to solution.

Molecular dynamics (MD) simulations have become indispensable for addressing this gap. Simulations of entire crystal unit cells show that proteins retain substantial internal flexibility.^31–37^ However, MD currently accesses only motions up to the µs regime, leaving slower conformational changes, which are often central to biological function, difficult to capture.

Nuclear magnetic resonance (NMR) spectroscopy is a unique technique to probe protein dynamics in various environments, including solutions, crystals, and complexes. Direct comparisons of solution-state NMR with magic-angle spinning (MAS) NMR of protein crystals yield valuable insights into the modulation of motions by the crystal lattice. Spin-relaxation experiments targeted towards backbone-amide or methyl sites showed that fast (sub-µs) local motions are largely preserved in crystals, except in loops engaged in packing contacts.^23,38,39^ By contrast, there is evidence that slower motions on the µs–ms timescale, which usually involve the concerted rearrangement of larger structural elements, are more strongly perturbed: a well-characterized case is the reshuffling of a β-turn in ubiquitin. Although this process persists in crystals, it is slowed by more than an order of magnitude, and different space groups alter the populations to a varying degree.^40,41^ Until now, rarer concerted motions involving even larger parts of a protein have not been studied by NMR methods.

Aromatic ring flips provide particularly sensitive probes of collective dynamics. Since the 1970s, NMR experiments have shown that phenylalanine and tyrosine rings undergo flips around the C^γ^ – C^ζ^ axis (the *χ*_2_ angle; Figure 1A).^42–44^ While surface-exposed rings often flip freely, buried rings require larger, more cooperative motions of the protein, sometimes referred to as “breathing motions”.^45,46^ Because the two interchanging states are structurally indistinguishable, crystallography cannot detect these flips. In contrast, they are easily observable by NMR, since the ^1^H and ^13^C spins at the δ and ε positions differ in chemical shift, producing line broadening or exchange cross-peaks depending on the timescale (Figure 1B). The introduction of α-ketoacid precursors (Figure 1C) for site-specific labeling of the (CH)^ε^ atoms in phenylalanine and tyrosine has greatly enhanced the precision of such studies.^47^

**Figure 1.**
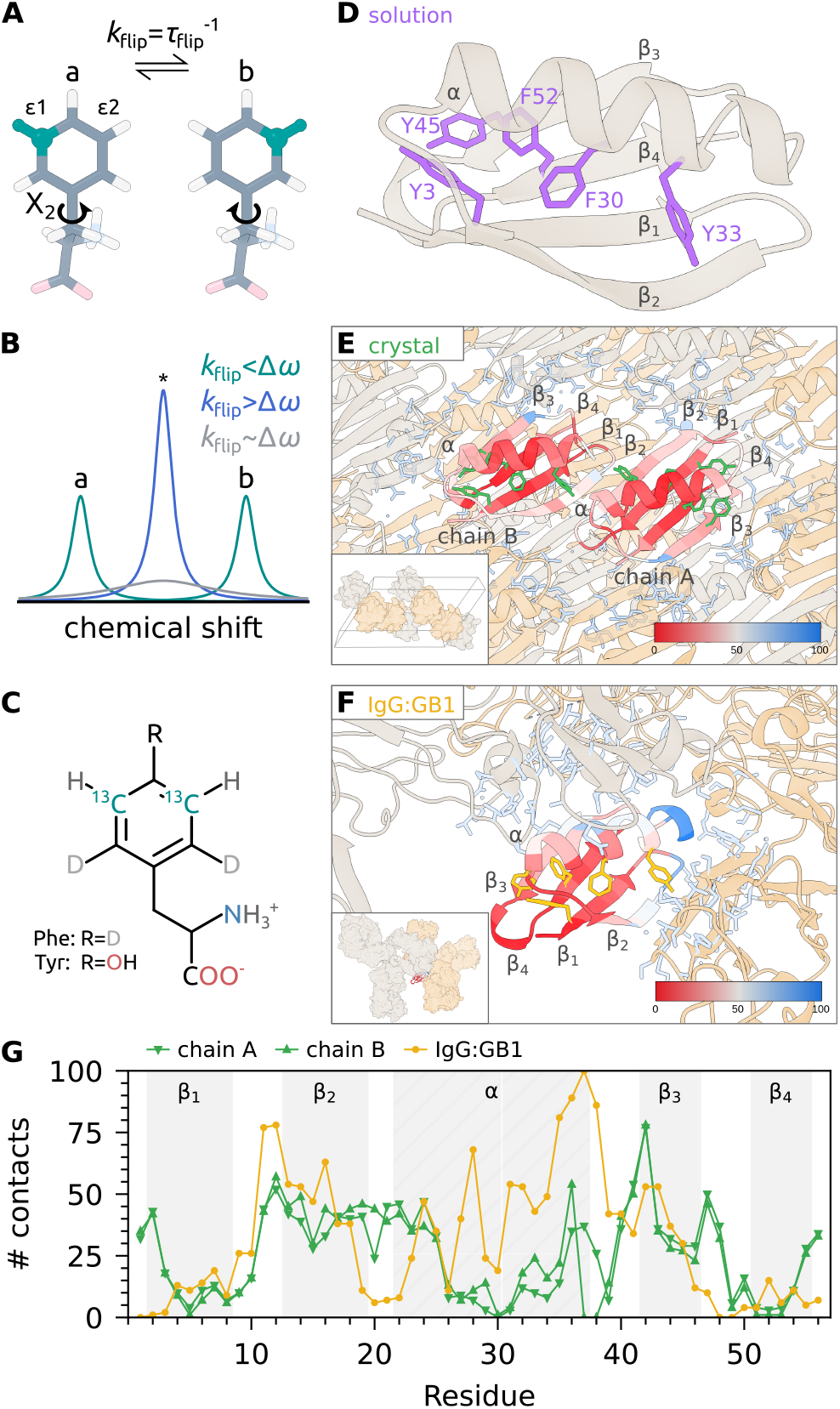
Specific isotope labeling of aromatic residues in different forms of GB1. (A) Schematic representation of a ring flip. For a ring flip, the *χ*_2_ angle around the C^*β*^ – C^*γ*^ bond rotates 180° with a flip rate *k*_flip_. (CH)^*ε*1^ and (CH)^*ε*2^ exchange their position in the ring, creating states a and b. (B) Schematic NMR spectra resulting from ring flips of (CH)^*ε*^-labeled Phe or Tyr residues at different timescales. If the ring flips on a timescale slower than the chemical shift difference Δ*ω* of sites a and b, the two signals are resolved (cyan). If *k*_flip_ *>* Δ*ω*, one averaged signal can be observed (blue). For intermediate exchange timescales where *k*_flip_ ~ Δ*ω*, the signal is broadened (gray). (C) α-Ketoacid precursor for 3,5-^13^CH labeling of the aromatic ring of Phe and Tyr residues. ^47^ (D) Structure of GB1_QDD_ in cartoon representation. Phe and Tyr residues are shown as purple sticks. (E) Surrounding of GB1_QDD_ in the crystal lattice. The asymmetric unit contains two molecules, denoted as chains A and B. The color gradient of the backbone encodes the number of atoms within 10 Å of the respective C^*α*^ atom. These atoms are shown as light blue sticks. The insert shows one unit cell with A and B molecules in different colors. (F) Surrounding of GB1_QDD_ in the IgG:GB1 complex. The insert shows the whole complex with two IgG and one GB1 molecule. The structure of the complex was modeled as described in. ^51^ (G) Number of atoms within 10 Å of C^*α*^ atoms in chains A and B of the crystal and the IgG:GB1 complex. Secondary structure elements are indicated as in the structure representations in (D)-(F).

Here, we use aromatic ring flips as sitespecific probes to assess how protein–protein interactions modify dynamics. We focus on the immunoglobulin-binding domain of protein G (GB1), studied in three contexts: in solution (Figure 1D), in the crystalline state (Figure 1E), and in complex with its natural binding partner, the immunoglobulin G (IgG; Figure 1F). The commonly studied T2Q mutant of GB1 (referred to here as wild-type), and the stabilized triple-mutant GB1_QDD_ (T2Q, N8D, N37D^48^) have long served as NMR model systems for structural and dynamical investigations. GB1_T2Q_ has been characterized extensively by solution and MAS NMR, and multiple crystal structures are available. GB1_QDD_, while structurally uncharacterized, has been studied in solution by NMR over a wide range of temperatures and hydrostatic pressures, providing a detailed baseline for comparison.^49,50^ In GB1, two phenylalanines (F30, F52) and two tyrosines (Y3, Y45) form a hydrophobic aromatic cluster, while Y33 is exposed to the solvent (Figure 1D). Backbone dynamics of GB1 in complex with IgG has been studied by MAS NMR,^51,52^ showing that while ps-ns motions are similar to the crystalline preparation, GB1 experiences an overall rocking motion on a µs time scale.^53^

Our MAS NMR measurements show that crystal packing slows ring flips of buried residues by approximately three orders of magnitude. To place these results in a structural context, we determined the crystal structure of GB1_QDD_. Multimicrosecond MD simulations of the protein in solution and as a crystal confirm the free-energy barriers obtained from experimental data and hint towards structural origins of the elevated barrier in crystals. Remarkably, in the IgG complex, the same ring flips are much faster than in crystals, highlighting how the precise nature of intermolecular contacts reshapes the underlying free-energy landscape.

## Results

### The packing of GB1_QDD_ in crystals and the IgG:GB1_QDD_ complex

Precise atomic-resolution structures are important for the mechanistic interpretation of protein dynamics. To our knowledge, crystallization of GB1_QDD_ has not been previously reported. For GB1_T2Q_, a commonly used crystallization protocol yields crystals in the P3_2_21 space group, suitable for solid-state NMR studies.^54^ However, under these conditions, crystallization of GB1_QDD_ reproducibly failed. We attribute this to the additional N8D and N37D mutations, which are surface-exposed and likely disrupt the packing interactions required for formation of the P3_2_21 lattice. Instead, we reliably obtained GB1_QDD_ crystals in the C121 space group using a dialysis-based method previously reported for other GB1 variants.^55^ These crystals were suitable for MAS NMR and X-ray diffraction (XRD), depending on the duration of crystallization (Figures S1, S2, and Supplementary Methods). Single-crystal XRD data were collected at a resolution of 1.08 Å (Table S1), revealing an asymmetric unit with two molecules in distinct local environments.

We compared the intermolecular packing in the GB1_QDD_ crystal with that in the IgG:GB1_QDD_ complex, reasoning that differences in packing could influence aromatic ring dynamics. As a quantitative measure of packing, we determined the number of non-hydrogen atoms within 10 Å of each C^α^ atom for chains A and B in the crystal (Figure 1E) and in a model of the IgG:GB1 complex based on crystal structures of GB1 or GB3 in complex with fragments of IgG^56,57^ (Figure 1F). The two crystallographic chains exhibit very similar packing densities (Figure 1G, green). In contrast, the distribution of contacts in the crystal differs markedly from that in the IgG complex (yellow). Although the overall extent of intermolecular contacts is comparable, their differing distributions provide an opportunity to dissect how distinct packing modes modulate protein dynamics.

### MAS NMR of GB1_QDD_ crystals

Equipped with this structural knowledge, we prepared crystals with two different isotope-labeling schemes by heterologous expression in *E. coli*. For studies of aromatic residues, we produced deuterated protein with isolated ^13^C^1^H pairs at the ε positions of phenylalanine and tyrosine rings using a metabolic precursor (Figure 1C; see Methods in the Supporting Information),^47^ hereinafter called F^*ϵ*^/Y^*ϵ*^-GB1. Combined with ^1^H-detection and fast MAS (40–55 kHz), this labeling strategy results in high-resolution spectra, with simple two-spin systems that are well suited for quantitative analysis of dynamic parameters. An additional labeling scheme, used for backbone assignment, consisted of uniform ^2^H, ^13^C, and ^15^N labeling. (In all cases, exchangeable hydrogen sites were protonated.)

Because the crystals used for NMR and X-ray diffraction, though grown under similar conditions, differed in size, we sought to verify that they crystallized with the same packing. To this end, we performed powder XRD-like measurements on wet microcrystalline samples closely resembling those used for NMR, and compared the resulting patterns with the single-crystal XRD structure. The microcrystalline data could be indexed with the same unit cell and space group (Figure S3). We therefore conclude that the crystals used for both XRD and NMR represent the same phase.

As GB1_QDD_ crystals have not been studied previously by MAS NMR, we performed experiments to assign the H^N^, N^H^, C’, C^α^ and C^β^ resonances using a set of three-dimensional experiments (see Supplementary Information). The high quality of the data allowed automatic assignments for most of the protein (Table S3, Figures S5 and S6, and Figure 2A). We observed two distinct sets of peaks for four residues (Q2, Y3, T18, V21), which we ascribe to the two non-equivalent molecules in the unit cell. A Na^+^-ion bound in the vicinity of one of the two chains may explain the differences.

**Figure 2.**
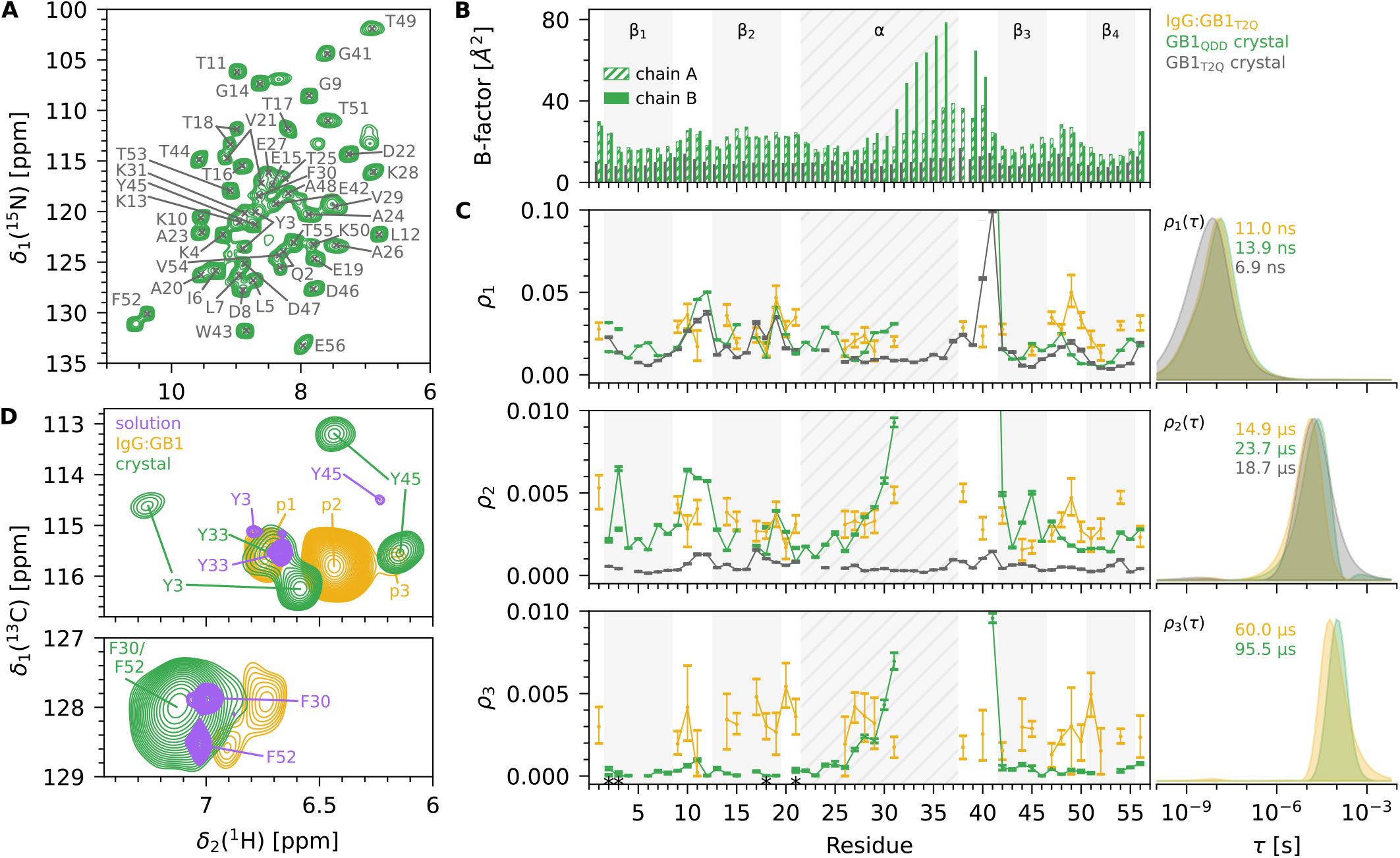
MAS NMR of different forms of GB1. (A) Dipolar-based ^1^H – ^15^N NMR spectra of crystalline GB1_QDD_ with backbone amide assignment. (B) B-factors of the GB1_T2Q_ (PDB 2QMT) and the GB1_QDD_ (PDB 9I2I) crystal with chains A and B. (C) Detectors analysis of backbone-amide ^15^N spin-relaxation data for the IgG:GB1_T2Q_ complex (yellow), ^53,58,59^ GB1_QDD_ microcrystals (green) and GB1_T2Q_ microcrystals (gray). ^58,60^ The panels on the left show residue-wise responses to the corresponding detector sensitivities *ρ*_1*−*3_(*τ*) on the right. *ρ*_1*−*3_(*τ*) differ slightly for each sample state, as the recorded relaxation rates were different; the correlation times corresponding to the maximum of the detector sensitivities for each sample are specified in each panel. (D) Tyrosine (upper panel) and phenylalanine (lower panel) region of ^1^H – ^13^C solution-state or dipolar-based MAS NMR spectra of F^*ϵ*^/Y^*ϵ*^-GB1 in solution (purple), a complex with IgG (yellow) and a microcrystal (green). Spectra of the complex and the crystal were recorded at MAS frequencies of 39 and 30 kHz, respectively.

Interestingly, no signals of residues Q32-D40 have been observed in the spectra. This part comprises the C-terminal half of the α-helix and the loop preceding the β3-strand. We ascribe this to dynamics on timescales that lead to very fast relaxation (further investigated below), as there are no unassigned resonances in the spectra and the crystallographic B-factors are high in this region (Figure 2B). Residues D37 and G38 in chain B of the crystal structure could not be modeled due to severe blurring of the electron density. In the previously reported MAS NMR studies of GB1_T2Q_ crystals and IgG-bound GB1_T2Q_, residues Q32-D40 were observable in NMR spectra, and the B-factors in the GB1_T2Q_ crystal are relatively low and uniform throughout the structure. These findings suggest crystal-packing-dependent differences in the dynamics of this region of the protein.

To quantitatively characterize the dynamics of the newly reported crystal, we performed ^15^N spin-relaxation experiments (longitudinal *R*_1_ relaxation and rotating-frame *R*_1*ρ*_ relaxation at six different spin-lock frequencies *ν*_SL_), and analyzed them using the detectors approach (Figure 2C, green; Figures S11 and S12).^53,58–61^ For comparison, we show previously published data of the GB1_T2Q_ crystal (gray)^58,60^ and the IgG:GB1_T2Q_ complex^53,58,59^ (yellow). The detectors report on the amount of motion within certain time windows, so-called detector sensitivities *ρ*_i_(*τ*) (Figure 2C, right). The position of these windows is dictated by the time scales of motion the measured relaxation rates are sensitive to. The sensitivities used herein cover a few ns to hundreds of ns (*ρ*_1_), several µs to hundreds of µs (*ρ*_2_), or tens of µs to a few ms (*ρ*_3_). The responses *ρ*_*i*_ (Figure 2C, left) report the motional amplitude of each residue in the respective sensitivity *ρ*_i_(*τ*).

The most striking feature is the elevated µs– ms motion for residues 28-31 and 41 (seen as a sharp increase of *ρ*_2_ and *ρ*_3_). As motion on this time scale also leads to strong line broadening and rapid signal loss,^62^ these data support the assumption that residues Q32-D40 are unobservable because of enhanced microsecond motions of the C-terminal half of the α-helix and the adjacent loop. Interestingly, this motion is not observed in GB1_T2Q_ crystals (Figure 2C, gray data set) and has not been reported in residual-dipolar coupling analyses in solution.^63^ Indirect evidence for local helical fraying has been presented based on the pH dependency of solution-NMR spectra.^64,65^ In addition to this local µs motion, we note that there is also an overall increase of *ρ*_1_ and *ρ*_2_ in GB1_QDD_ and IgG:GB1, compared to the previous crystal form. For IgG:GB1, this finding has been ascribed to the overall rocking motion of the protein,^52^ which can also be found in crystals.^35^ Fast motions (*ρ*_1_) are similar in the three forms, presumably due to their local nature.

### Different NMR signatures of aromatics in three different states of GB1_**QDD**_

We obtained high-resolution dipolar-based ^1^H – ^13^C 2D MAS NMR spectra of crystalline GB1_QDD_ and the IgG:GB1_QDD_ complex and compared them to solution-state NMR spectra of GB1_QDD_ (Figure 2D). The spectral signatures directly show that the ring flip dynamics differ in these samples, as further investigated below. To interpret these differences, residue-specific assignments of the signals are needed. We performed experiments that exploit the spatial proximity of the (^13^C^1^H)^ε^ sites to the surrounding amide signals, for which the assignments have been obtained (see above). Three-dimensional Radio Frequency-Driven Recoupling (RFDR)^66^ experiments allowed us to assign all tyrosine signals in the crystalline state (Figure S7); the signals from the two phenylalanine residues are not resolved and only a single broad peak was observed (Figure 2D, lower panel).

While this assignment approach worked well for crystals, low sensitivity of the IgG:GB1_QDD_ sample precluded residue-specific assignments of the complex. We nonetheless sought to understand if the different Tyr peaks are from individual residues, or if there are any peaks arising from slow ring flips. No cross-peaks were found in a ^13^C – ^13^C EXchange SpectroscopY (EXSY) experiment (Figure S10), indicating that each of the signals observed in the IgG:GB1_QDD_ spectrum originates from an individual residue, rather than from slow exchange. We therefore refer to the three Y^*ϵ*^ peaks of IgG:GB1 as p1, p2, and p3 in the following.

### Differences in aromatic spectra unambiguously show altered ring flip dynamics

The number of signals observed for a (^13^C^1^H)^ε^ site provides a direct, semi-quantitative view of the ring-flip rate constant *k*_flip_ (Figure 1B). For GB1_QDD_ in solution, each of the five labeled aromatic residues gives rise to a single averaged signal because *k*_flip_ *>* Δ*ω*, where Δ*ω* is the chemical shift difference between (CH)^ε1^ and (CH)^ε2^ (Figure 2D, purple). With typical values of Δ*ω*, this observation means that the ring flips occur on timescales of µs or faster. Previous relaxation-dispersion experiments allowed quantifying the rate constants:^49^ At 298 K, the ring flips all occur at a rate constant of a few tens of thousand per second (Y3: (13 700 ± 1600) s^−1^; Y45: (42 100 ± 21 700) s^−1^; F30: (29 600 ± 1500) s^−1^; F52: (34 400 ± 4500) s^−1^). The solvent-exposed Y33 flips too fast for quantification by solution NMR relaxation-dispersion experiments, implying the ring flips to occur at least at 100 000 s^−1^.

Similarly to the situation in solution, a single set of peaks is observed in the IgG:GB1 complex (Figure 2D, yellow).

In clear contrast to solution and the IgG:GB1_QDD_ complex, the spectrum of the crystalline sample exhibits a very different signature, namely two peaks for each Y3 and Y45 (Figure 2D, green). The peaks are split approximately equidistantly around the position found in solution. This finding strongly suggests that these two residues flip more slowly in the crystal than in solution, such that *k*_flip_ *<* Δ*ω*, and fast flips in solution average the signals to a single signal. The solvent-exposed Y33 features a single signal implying fast flips. The single broad peak in the phenylalanine region also stands in contrast to the sharp single lines observed in solution, and points to µs motion of the F30 and/or F52 in crystals.

### Crystal packing slows flips of buried tyrosine rings by three orders of magnitude

To quantify *k*_flip_ of Y3^crystal^ and Y45^crystal^, we performed ^1^H – ^13^C EXSY measurements (Figure 3).^67^ This type of experiment probes the respective buildup and decay of cross and diagonal peaks during an increasing exchange delay, *τ*_ex_. Diagonal peaks “a” and “b” are the signals of the two sites (CH)^ε1^ and (CH)^ε2^, while cross peaks “ab” and “ba” connect frequencies corresponding to the states that are interconverted by the ring flip (Figure 3A). Figure 3B shows the normalized intensities of the signals depending on the exchange delay. The extracted flip rates for Y3 and Y45 are (55.8 ± 2.2) s^−1^ and (18.37 ± 0.11) s^−1^ at 304 K, i.e., about three orders of magnitude slower than in solution.

**Figure 3.**
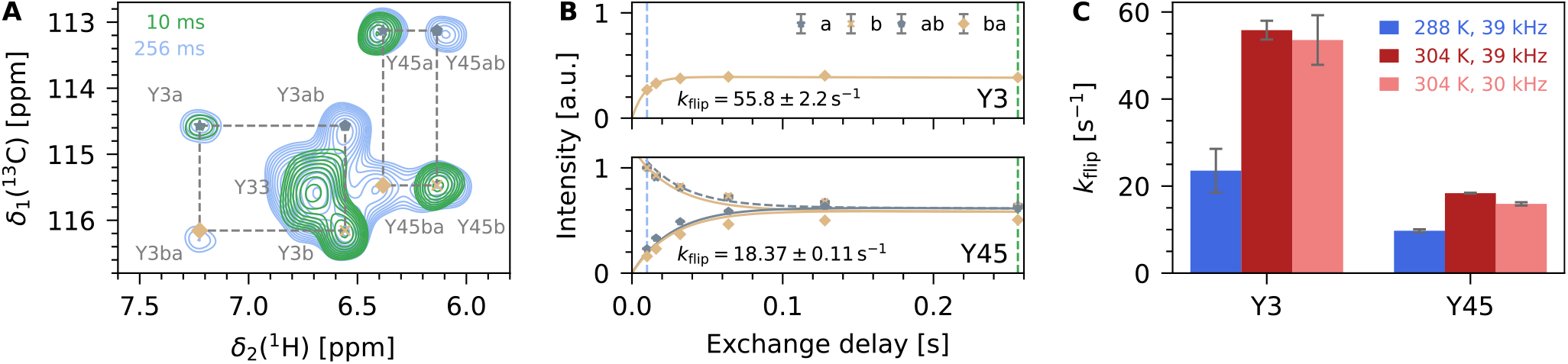
Determination of ring flip timescales of Y3 and Y45 in crystalline GB1_QDD_. (A) ^1^H – ^13^C EXSY MAS NMR spectrum of Y3 and Y45 in microcrystalline GB1 with 0 (green) and 256 ms (light blue) exchange delay recorded at a MAS frequency of 39 kHz. (B) EXSY buildup and decay curves for Y3 and Y45. For Y3, only the buildup of “ba” was fitted as the intensities of the other signals were strongly impacted by peak overlap with Y33. For Y45, we performed a combined analysis of all four peaks. ^69^ (C) Temperature- and MAS-dependence of *k*_flip_ for Y3 and Y45. The apparent rate constants are approximately doubled at 304 K compared to 288 K and are hardly dependent on the MAS frequency; in fact, the modest change is the inverse of what one would expect if the buildup arose due to spin diffusion.

To ensure that the observed EXSY cross-peaks are indeed due to ring flips, as opposed to magnetization transfer (spin diffusion), we performed control experiments (Figure 3C and S13). Magnetization transfer in MAS NMR – which would occur even in static systems^68^ – depends on the MAS frequency, whereas ring-flip dynamics are expected to depend on temperature. The observation that the buildup rate constants do not change with MAS frequency, but increase as the temperature is raised, allows us to conclude that the rate constants observed in the EXSY experiment report on ring-flip rate constants.

### Probing fast-flipping rings with dipolar couplings and spin relaxation

For sites with only a single signal, EXSY is not applicable. Instead, we measured dipolar order parameters and ^13^C spin relaxation experiments to gain insights into ring flip kinetics (Figure 4). The ^1^H – ^13^C dipolar order parameter *S* (defined as 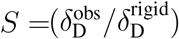, where 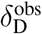 is the measured dipolar-coupling tensor anisotropy and 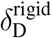 is the dipolar-coupling tensor anisotropy in the absence of motion), ranges from 1 for totally rigid sites to 0 for flexible disordered sites. It reports on amplitudes of motion on timescales up to hundreds of µs.^70^ The motional averaging due to a ring flip results in a theoretical value of *S* = 0.625;^71^ reductions below this value point to additional motions, such as the libration of the ring within each rotamer state, motion of the backbone, or the ring axis (i.e., the *χ*_1_ angle). We determined dipolar order parameters using Rotational-Echo DOuble-Resonance (REDOR) experiments (Figure 4A, S14, and S15). In the crystal, Y3 and Y45 have dipolar order parameters *S* of approximately 0.8, i.e., higher than the value expected for ring flips. This finding unambiguously confirms that Y3 and Y45 do not flip on timescales shorter than about 100 µs. In contrast, Y33 and the phenylalanines have low order parameters, indicating that these rings flip faster than roughly one-hundred µs.

**Figure 4.**
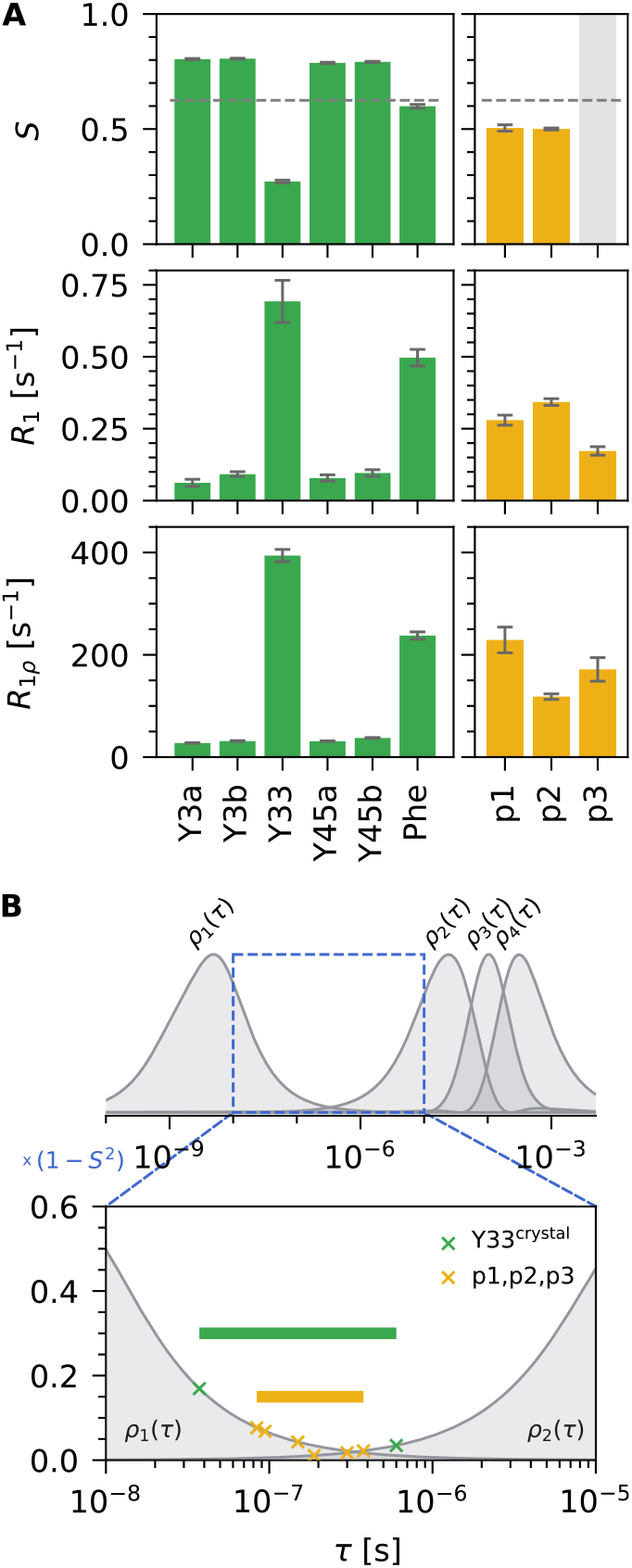
Determination of ring flip timescales of fast-flipping tyrosines in GB1. (A) Dipolar order parameters *S* (top), ^13^C^*ε*^ *R*_1_ (middle) and *R*_1*ρ*_ (bottom) relaxation rate constants of Tyr and Phe residues in crystalline GB1 (left) and in the IgG:GB1 complex (right). Measurements were performed at a MAS frequency of 39 kHz and *R*_1*ρ*_ experiments were measured with a spin-lock field strength *ν*_SL_ of 15 kHz. (B) Estimation of ring flip rates for tyrosines Y3^crystal^, p1, p2, and p3. The upper panel shows the detector sensitivities *ρ*_1*−*4_(*τ*) that resulted from a detectors analysis of six *R*_1*ρ*_ rate constants (*ν*_SL_ = 15, 25, 27.5, 30, 32.5 and 35 kHz) and *R*_1_. The dashed box indicates timescales between the maxima of *ρ*_1_ and *ρ*_2_ that our set of experiments is not sensitive to. The lower panel shows a close-up of that area with the sensitivities scaled by (1 *− S*^2^) with *S* = 0.625 being the theoretical order parameter of a ring flip. The crosses mark the intersection between the scaled sensitivities and the respective responses. The corresponding correlation times mark the possible range of timescales for the ring flips indicated by horizontal bars.

To quantify the ring-flip time scales more precisely, we used ^13^C spin relaxation experiments, in particular ^13^C *R*_1_ (sensitive to motions on the hundreds of picoseconds to tens of nanoseconds time scale) and *R*_1*ρ*_ (hundreds of ns to hundreds of µs. The markedly different relaxation behavior of Y33, compared to the slowly-flipping Y3 and Y45, immediately points to faster dynamics (Figure 4A). We used the detectors approach^58,61^ with seven experimental relaxation rate constants to determine the amplitude of motion within each of the four detector sensitivities *ρ*_1*−*4_ (Figure Figure 4B, upper panel, and Figure S16, S17, and S19). The theoretically expected response for a ring flip at a specific timescale allowed us to define the upper and lower bounds of the ring-flip rate constants for Y33^crystal^, which are 37 and 600 ns, respectively (Figure 4B lower panels, S8, and details in the methods section).

The broad Phe signal, ascribed to F30 and/or F52, allows us to obtain insight into the motion of these two residues located in the hydrophobic core. The order parameter (*<* 0.625) points to ring flips on a timescale shorter than hundreds of µs. The high *R*_1_ and *R*_1*ρ*_ relaxation rate constants are compatible with Phe rings flipping on a ns-µs timescale, which is in line with the large line width. Due to the overlap of the two phenylalanine signals, extracting site-specific information for F30 and F52 is not possible; therefore, we focus on the tyrosines in the following discussion.

### Ring flips in the IgG:GB1_QDD_ complex are faster than in crystals

The ^1^H – ^13^C 2D MAS NMR spectra indicate that the aromatic residues of GB1_QDD_ in complex with IgG have different ring-flip dynamics than those in the crystal lattice: the fact that there is only one peak per tyrosine points to fast (sub-ms) ring flips (Figure 2D). Low dipolar order parameters and elevated ^13^C *R*_1_ and *R*_1*ρ*_ relaxation rate constants support this view (Figure 4A and S18). Similarly to the quantification of Y33^crystal^ dynamics above, a detectors analysis of these relaxation data allowed us to estimate the range of time scales of the ring flips. The range lies between tens to hundreds of nanoseconds (Figure 4B and S19) and is similar for all three Tyr peaks (p1, p2, p3). Although we do not have residue-specific assignments, we can conclude that the Tyr ring flips in the IgG:GB1_QDD_ complex occur on the highns time scale, i.e., about five orders of magnitude faster than in the crystal. This finding implies that the intermolecular contacts in the complex favor states that allow ring flips. We speculate that inter-molecular contacts favor more extended states of GB1 by loosening the hydrophobic core. The absence of high-resolution structures of GB1 in complex with full-length IgG precludes the detailed analysis of the mechanisms.

### Reduced disorder of the transition state is the primary reason for slow flips in crystals

The ring flip correlation times *τ*_flip_ of the tyrosines in all three states of GB1_QDD_ are summarized in Figure 5A. We sought to characterize properties of the transition state of the ring flips by studying the temperature-dependence of *k*_flip_ of Y3 and Y45 (Figure 5B). While the absolute rate constants in solution and crystals are very different, the slopes of log *k* as a function of 1*/T* in the crystal are similar, albeit slightly flatter in crystals than in solution. This observation can only be explained if the activation enthalpy Δ*H*^*‡*,solution^ *≥* Δ*H*^*‡*,crystal^, as one can readily show from the Eyring equation (see Supporting Information, Figure S9). The large vertical offset of *k*_flip_ must then be due to differences in the activation entropy Δ*S*^*‡*,solution^ *>* Δ*S*^*‡*,crystal^. In other words, the transition states of the ring flips of Y3 and Y45 in crystals are less disordered compared to those in solution.

**Figure 5.**
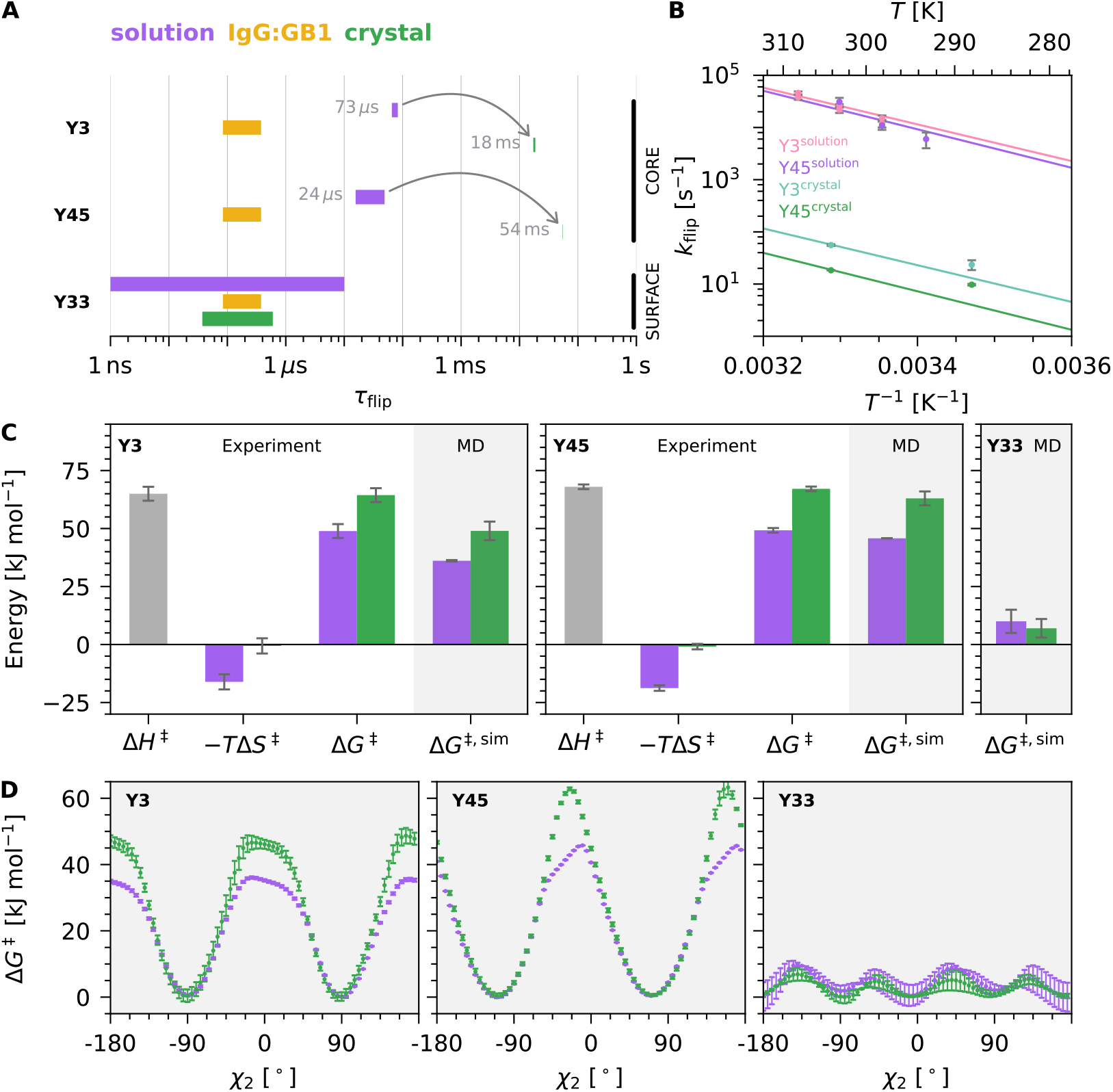
Thermodynamic parameters of tyrosine ring flips seen from experiment and simulations. (A) Overview of the correlation time constants of ring flips in the three sample states at 298 K (solution) or 304 K (crystal and complex). The bars indicate the range of the determined timescales, including uncertainties. Values for Y33^crystal^ (green) and the three tyrosines in the IgG complex (yellow) were obtained from the detectors analysis of ^13^C spin relaxation; due to lack of site-specific assignments, the maximum range of time constants that includes p1, p2, and p3 is shown here. Values for Y3^crystal^ and Y45^crystal^ were obtained from EXSY. Data for Y3^solution^ and Y45^solution^ come from ^13^C *R*_1*ρ*_ experiments; ^49^ the upper limit for Y33^solution^ flips is obtained from theoretical considerations regarding the limits of solution NMR relaxation dispersion measurements. (B) Temperature dependence of flip rate constants of Y3 and Y45 in solution ^49^ and crystals. The solid lines show a per-residue joint fit of solution and crystal data, constraining Δ*H*^*‡*,solution^ = Δ*H*^*‡*,crystal^. (C) The fitted Δ*H*^*‡*^ and *T* Δ*S*^*‡*^ from (B) were used to calculate activation free energies Δ*G*^*‡*^ at *T* = 298 K for Y3 and Y45. Δ*G*^*‡*,sim^ are the free energies from enhanced-sampling MD simulations (see panel D of this figure). The parameters are shown for solution (purple) and the crystal (green), except for Δ*H*^*‡*^ (gray), which was set to Δ*H*^*‡*,solution^ = Δ*H*^*‡*,crystal^ in the fit (see (C)). (D) Δ*G*^*‡*^ profiles for ring flips of Y3, Y33, and Y45 obtained from enhanced-sampling MD simulations of GB1 in solution (purple) or the crystal (green).

For a more quantitative analysis, we fitted this data with the Eyring equation. As the temperature range was limited for the MAS NMR measurements (by the available cooling power against the MAS-induced heating and sample stability), we performed a joint fit of the rate constants from solution and crystal with the restriction that Δ*H*^*‡*,solution^ = Δ*H*^*‡*,crystal^ for each residue. (Additionally, we also tested Δ*H*^*‡*,solution^ *>* Δ*H*^*‡*,crystal^, which results in a very similar overall conclusion.) Figure 5C shows the fitted parameters Δ*H*^*‡*^ and *T* Δ*S*^*‡*^ at *T* = 298 K as well as the resulting Gibbs free energy of activation Δ*G*^*‡*^ at *T* = 298 K. The large difference in activation entropy, Δ*S*^*‡*,solution^ *≥* Δ*S*^*‡*,crystal^, reveals that the flips of Y3 and Y45 are slowed down due to a higher entropic barrier in the crystal compared to solution. A lower degree of disorder in the crystal is expected as the crystal lattice imposes constraints on the protein. Comparing the activation energies between solution and crystal results in ΔΔ*G*^*‡*^ = (14 *±* 9) kJ mol^*−*1^ for Y3 and (17 ± 3) kJ mol^−1^ for Y45, a value that is useful to benchmark simulations shown below.

We also measured ^13^C *R*_1*ρ*_ spin relaxation at a lower temperature and analyzed them with the detectors approach to obtain temperature-dependent *k*_flip_ of Y33^crystal^ (Figure S16 and S19). However, the data did not show temperature dependence within experimental error. This is presumably due to the limited temperature range available for MAS measurements and the fact that the detectors approach only allowed us to determine an upper and lower bound for *k*_flip_ and not an exact value. We therefore refrained from further analyzing the temperature-dependent data for Y33.

### Enhanced-sampling molecular dynamics simulations quantitatively recapitulate experimental ring-flip kinetics

To gain mechanistic insights into the ring flips, we performed enhanced-sampling moleculardynamics simulations.^72^ MD simulations provide atomic-level information on protein dynamics in solution and crystals^31–36^ and are highly complementary to experimental methods, which generally do not provide information about each atom. With increasing computational power, simulations of large crystal lattices, comprising multiple copies of the unit cell, and extending over the µs-timescale, have become feasible.^73^ However, the question at hand remains challenging, given that the ring flips occur on timescales spanning from tens of µs in solution to tens of ms in the crystal. Such rare events are not amenable to current allatom equilibrium MD simulations carried out on non-specialized computer architectures.^74^

Enhanced-sampling MD simulations^72^ allowed us to accelerate the exploration of the conformational space along the *χ*_2_ angle, utilizing the well-tempered metadynamics-extended adaptive biasing force (WTM-eABF) algorithm^75^ in its multiple-walker variant,^76^ as implemented in NAMD^77^ and Colvars.^78^ Details of the method and the performed simulations are provided in the Supporting Information. As shown in Figure 5D, in both aqueous solution and crystal environments, our µs-timescale simulations show that the free-energy barriers against the ring flips of the buried Y3 and Y45 are substantially higher than the one of the more exposed Y33. This result is in quantitative agreement with our experimental observations (Figure 5C). Importantly, the absolute values of Δ*G*^*‡*^ agree well with those determined experimentally. This good quantitative agreement of the free-energy barrier heights between experiments and MD simulations provides a solid foundation for a mechanistic understanding of the ring dynamics.

### Ring dynamics involve a complex network of interactions, but not breathing motions

In search for mechanisms underlying the ring flips, we investigated potential temporal correlations between ring flips of Y45 (i.e., *χ*_2_) and fluctuations in other structural parameters, such as distances and angles. We initially hypothesized that flips of this buried aromatic ring might be coupled to global “breathing” motions. While this term lacks a precise definition, we analyzed fluctuations in the compactness of the hydrophobic core by monitoring several distances across the core of the protein such as the distance between the C^α^ atoms of residues 45 (located in β-strand 3) and 23 (located in the α-helix). No detectable correlation was observed between fluctuations in the Y45 ring-axis orientation (*χ*_1_) or ring-flip state (*χ*_2_) and fluctuations of these long-range distances (Figure S21 and S22). This finding suggests that ring flips are not directly associated with a simple expansion or contraction of the hydrophobic core.

In the simulation of the GB1 monomer in solution, *χ*_2_ alternates between two well-defined states (approximately 50° and −130°, respectively). The ring axis, defined by the *χ*_1_ angle, predominantly remains in a single conformation and exhibits only rare, short-lived excursions of roughly 50° from this dominant state (Figure S20). There is no apparent temporal correlation between fluctuations in *χ*_1_ and *χ*_2_.

In the crystal lattice, the situation is more complex. Here, *χ*_2_ not only fluctuates between two symmetry-equivalent positions separated by 180°, but also populates intermediate states. Furthermore, excursions of the ring axis (from *χ*_1_ *≈* −25° to 20°) are longer-lived and often – albeit not always – coincide with transitions in *χ*_2_ (Figure 6G). In the two major *χ*_1_ states, the ring is rotated more or less “outward” (Figure 6A, C, E and B, D, F, respectively). These states are also associated with significant rearrangements in the intramolecular interactions. Of particular relevance are interactions involving the hydroxyl hydrogen of Y45, the carboxyl group of D47, and the positively charged ammonium moiety of K50, which differ substantially between the two conformations. However, as these interactions are intramolecular, they may also occur in solution, and, thus, cannot explain the observed deceleration of ring flips in the crystal.

**Figure 6.**
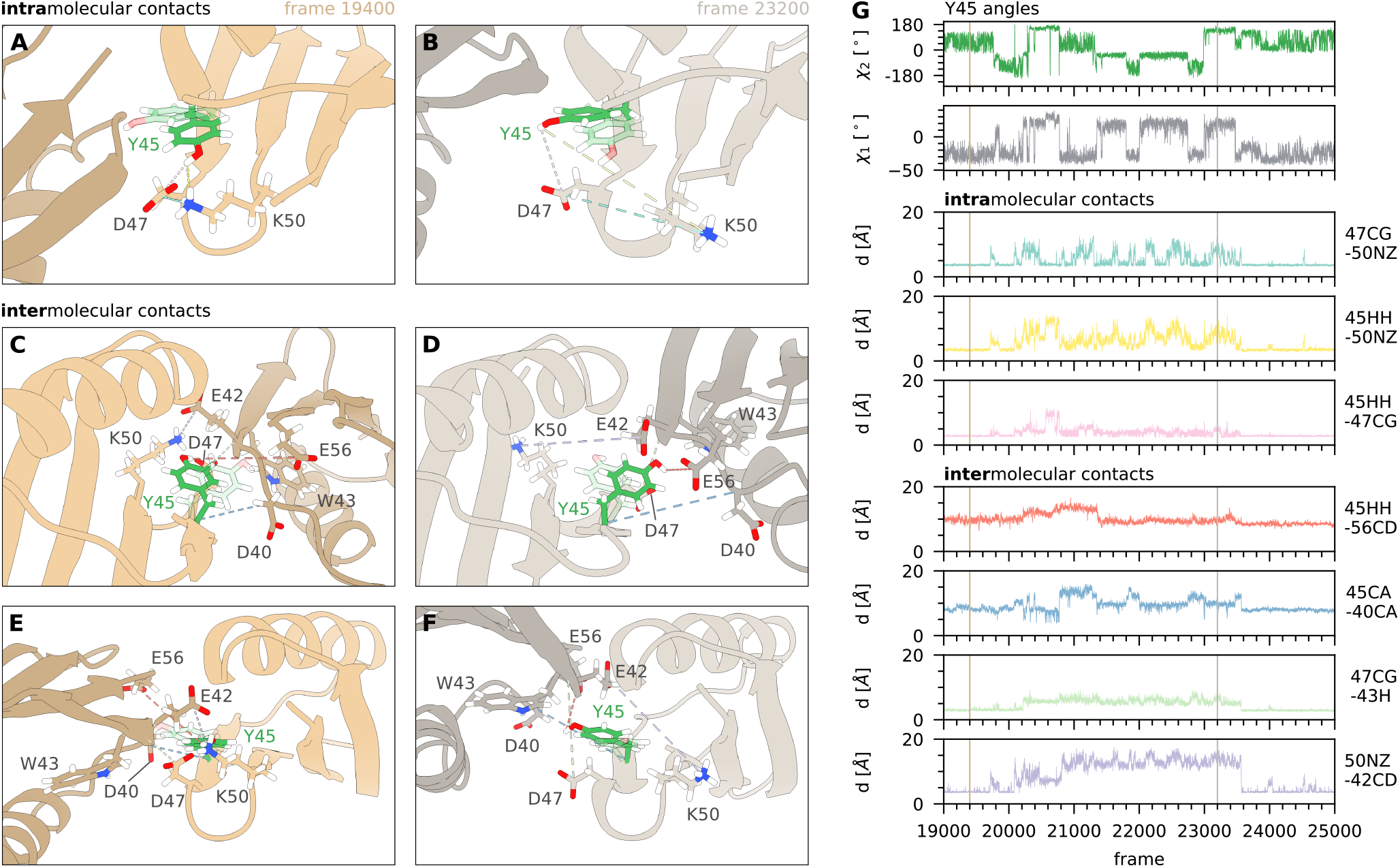
Structural insights into the deceleration of aromatic ring flips of Y45 in crystals compared to solution from MD simulations. (A-F) Close-up of Y45 and residues interacting with it intermolecularly (A-B) or in a neighboring molecule (C-F). (G) Time traces of the dihedral angles of the side chain of Y45, and relevant intra- and inter-molecular distances. The vertical lines indicate the frames corresponding to the structures shown in panels A-F. The full time trace, comprising over 56000 frames, as well as data of the simulation of the monomer (in solution), are shown in Figures S20 and S21.

We identified an additional set of intermolecular contacts that may offer mechanistic insight into the influence of the crystal lattice on ring flips. These contacts involve residues D40, E42, W43, and E56 from a neighboring protein subunit in the lattice (Figure 6C–G). Fluctuations in the distances between these residues and the hydroxyl group of Y45 – or its nearby interaction partners – tend to coincide with fluctuations in *χ*_1_ and/or *χ*_2_ of Y45. Since these contacts are necessarily absent in solution, we propose that they contribute to the increased free-energy barrier for ring flips observed in the crystal environment.

Overall, MD simulations indicate that ring dynamics are governed by a complex network of interactions. Rather than pointing to “breathing” motions in the sense of a physical expansion of the protein, we find the ring dynamics (both flips and ring-axis reorientation) to depend on a subtle interplay of intra- and intermolecular contacts.

## Discussion

Proteins function by sampling a spectrum of conformations. Consequently, they operate within a delicate balance of interactions, and their three-dimensional structures are typically only marginally stable. Even GB1_QDD_, considered a highly stable protein, has a free energy of unfolding (Δ*G*^0^ *≈* 20 kJ mol^−1^ ^48^) that corresponds to the energy of just a few hydrogen bonds. It is therefore not surprising that additional interactions with the environment can significantly reshape the protein’s free-energy landscape.

In this study, we used aromatic ring flips as sensitive reporters to investigate how interactions of GB1 with either neighboring molecules in the crystal or with a large binding partner (the IgG antibody) influence its dynamics compared to the protein in solution. Ring-flip dynamics are particularly informative because they probe the intricate interaction network within the hydrophobic core and reflect rare, cooperative motions involving multiple residues.

We observed a spectrum of behaviors for the tyrosine ring flips in GB1. Y33, located on the protein surface, appears largely unaffected by its environment. It undergoes rapid ring flips on the nanosecond timescale in all three examined states. This finding aligns with a recent study of phenylalanine flips in three different crystal lattices of ubiquitin.^79^ Specifically, a surface-exposed phenylalanine in ubiquitin (F4), lacking direct crystal contacts, exhibits consistent ring-flip timescales in all crystal forms (10–20 ns), mirroring the behavior of Y33 in GB1.

In sharp contrast, the ring flips of Y3 and Y45 in GB1_QDD_ are slowed by approximately three orders of magnitude. This is mechanistically attributed to a less disordered transition state compared to that in solution, leading to the entropic penalty revealed by our thermodynamic analysis (Figure 5C). The reduced capacity of GB1 to sample conformations that enable ring flips likely stems from contacts with neighboring molecules across the protein surface (Figure 1E). The network of contacts revealed in our MD simulations (Figure 6C-F) may constitute such a framework where disorder is reduced by defined interactions. Surprisingly, in the IgG complex, the ring flips of Y3 and Y45 are accelerated and even faster than in solution. We hypothesize that interactions between GB1_QDD_ and IgG stabilize conformations that favor ring flipping. This may be facilitated by the fact that GB1-IgG contacts occur primarily on the side of the protein opposite Y3 and Y45 (Figure 1F).

The phenylalanine residues in the aromatic cluster (F30, F52) display markedly different (much faster) ring-flip dynamics in the crystal compared to Y3 and Y45 (Figure 4A). This observation directly shows that in crystals the mechanisms enabling F30 and F52 flips must differ from those governing tyrosine rotations and that, therefore, ring flips in crystalline GB1_QDD_ cannot be described by a global “breathing motion”.

In solution, by contrast, all aromatics in the core (Y3, Y34, F30, F52) flip with a similar rate constant in the range of a few tens of thousands per second. This similarity has been interpreted as a collective motion of the aromatic cluster, such as a global “breathing” motion, that allocates sufficient space for the rings to flip. It is tempting to imply that in solution, the aromatic rings in the core flip via some sort of collective “breathing motion”, while in crystals this motion is quenched. But in fact, the situation may be more complex also in solution: high-pressure NMR measurements have shown that ring flips can proceed without an increase in the total protein volume at the transition state (Δ*V* ^*‡*^).^50^ This implies that there is not a single process that allows for rings to flip, but that alternative pathways exist that allow rings to flip also in solution via more “local breathing” motions/local rearrangements without a change in net volume^50^).

Recent data support that ring-flip dynamics may not be governed by a single global mode: Lu *et al*. used fluorine-labeled tyrosines in conjunction with solution-state ^19^F NMR to study ring flips in GB1_T2Q_ in solution, crowded environments, and inside cells.^80^ As fluorocarbons are more hydrophobic than hydrocarbons, fluorinated tyrosine rings are expected to engage in stronger local interactions, resulting in higher energy barriers for ring flipping. Consistent with this, the ring-flip rate constant of fluorinated Y45 is approximately 3–4 times lower than its non-fluorinated counterpart (9100 s^−1^ vs. (42 000 ± 22 000) s^−1^ at 298 K). However, flips of fluorinated Y3 are much faster than those of fluorinated Y45 (60 s^−1^ vs. 9100 s^−1^), showing that Y3 and Y45 do not flip synchronously in F-Tyr-labeled GB1_T2Q_. While this behavior may be due to the unique properties of fluorinated tyrosines, it underscores that timescales of aromatic ring flips are governed by multiple factors that likely involve a combination of local and collective motions.

Our data highlight that protein-protein interactions add to this complexity of molecular mechanisms governing ring flips: intermolecular contacts can slow ring flips down (as seen for Y3 and Y45 in crystals), but they may also help to stabilize states that facilitate ring flips, as evidenced by the accelerated flips in the IgG:GB1 complex.

Our findings may also have relevance for dynamics in the crowded cellular environment where protein-protein interactions are ubiquitous; in-cell NMR studies have shown that quinary interactions can promote partially unfolded states,^81^ including the case of GB1.^82,83^

In conclusion, our study demonstrates that intermolecular contacts can profoundly influence the timescale of internal protein dynamics. While the precise impact of the crystal lattice or complex certainly depends on the specific protein, our findings underscore that the protein free-energy landscape is complex and highly dependent on the surrounding environment.

## Acknowledgements

We thank Nikolai R. Skrynnikov and Olga O. Lebedenko (St. Petersburg) for insightful discussions and for performing exploratory MD simulations. We are grateful to Tobias Schubeis (Lyon) for advice with GB1 crystallization, Rebecca Schmid for initial crystallization trials, and Daniel Balazs (ISTA) for the powder XRD data collection. We thank Sebastian Falkner for assistance with constructing the structural model of the IgG:GB1 complex. This research was supported by the Scientific Service Units (SSU) of Institute of Science and Technology Austria (ISTA) through resources provided by the Nuclear Magnetic Resonance and the Lab Support Facilities. We thank Petra Rovó and Margarita Valhondo Falcón for excellent support of the NMR facility. Lea M. Becker is recipient of a DOC fellowship of the Austrian Academy of Sciences at the Institute of Science and Technology Austria (grant no. PR10660EAW01). Christophe Chipot acknowledges the European Research Council (grant project 101097272 “MilliInMicro”) and the Métropole du Grand Nancy (grant project “ARC”).

## Supporting Information

The Supporting Information file contains supplementary text, figures and tables covering details about protein production, crystallization and NMR sample preparation, experimental details about crystallographic data collection and structure determination, experimental details about NMR pulse sequences and acquisition parameters, backbone assignment of GB1_QDD_, including figures showing spectral assignments and tables of resonance frequencies, details about analyses of relaxation rate constants and dipolar-coupling measurements, including figures, details about MD simulations.

