## Supporting Information for "Aromatic Ring Flips Reveal Reshaping of Protein Dynamics in Crystals and Complexes"

[b] Haohao Fu
Tianjin Key Laboratory of Biosensing and Molecular Recognition, Research Center for Analytical Science, Frontiers Science Center for New Organic Matter, College of Chemistry, Nankai University, Tianjin 300071, China

[c] Matthias Dreydoppel, Ulrich Weininger
Institute of Physics, Biophysics, Martin-Luther-University Halle-Wittenberg, D-06120Halle (Saale), Germany

[d] Sylvain Engilberge*
European Synchrotron Radiation Facility, Grenoble Cedex 9 38043, France, and Univ. Grenoble Alpes, CNRS, CEA, Institut de Biologie Structurale, Grenoble Cedex 9 38044, France.


[e] Christophe Chipot*
Laboratoire International Associé CNRS and University of Illinois at Urbana-Champaign, LPCT, UMR 7019 Université de Lorraine CNRS, Vandoeuvre-les-Nancy F 54500, France, and Department of Physics, University of Illinois at Urbana-Champaign, Urbana, Illinois 61801, United States, and Department of Biochemistry and Molecular Biology, The University of Chicago, Chicago, Illinois 60637, United States.


**2 Supplementary Figures and Tables** **16**

### 1 Experimental

#### 1.1 Sample preparation

##### Protein production and purification

GB1 was produced either (1) without isotope labeling for structure determination, (2) with uniform $^{2}H$, $^{13}C$ and $^{15}N$ labeling for backbone assignment, or (3) with uniform $^{2}H$ and $^{15}N$ and site-specific $^{1}H-^{13}C$ labeling at the $(CH)^{\epsilon}$-positions of Phe and Tyr residues for studying aromatic ring dynamics. The M9 minimal medium (M9) for isotopic labeling contained: 5.25 g $L^{-1}$ $Na_2HPO_4$, 3 g $L^{-1}$ $KH_2PO_4$, 0.5 g $L^{-1}$ NaCl, 1 g $L^{-1}$ $NH_4Cl$, 2 g $L^{-1}$ D-Glucose, 1 mM $MgSO_4$, 0.1 mM $CaCl_2$, 0.1 mM $MnCl_2$, 0.05 mM $ZnSO_4$, 0.1 mM $FeCl_3$, 1 mg pyridoxine, 1 mg biotine, 1 mg D-pantothenic acid hemicalcium, 1 mg folic acid, 1 mg choline chloride, 1 mg niacinamide, 0.1 mg riboflavin, 5 mg thiamine hydrochloride.

Transformation of the plasmid containing the GB1 gene with the mutations T2Q, N8D, and N37D (*1*) into *E. coli* BL21(DE3) was achieved by a standard heat shock protocol and plated on LB agar containing 100 µg $mL^{-1}$ ampicillin (Amp). In the following, all shaking steps are performed at 37 °C and 200 rpm, all media formulations contain 100 µg $mL^{-1}$ Amp and cultures are inoculated such that the optical density at 600 nm ($OD_{600}$) is 0.2 unless stated otherwise.

For (1), a single colony was used to inoculate an overnight culture of LB medium. The next day, fresh LB medium was inoculated and shaken until it reached an $OD_{600}$ of 0.6 at which point expression was induced with 1 mM isopropyl-β-D-thiogalactopyranosid (IPTG). After shaking for 4 h, cells were harvested by centrifugation for 15 min at 4 °C and 5500 × g. The pellet was either frozen at −20 °C or used for protein purification immediately.

For (2), the initial preculture in LB was followed by three precultures in M9 with 0, 50 and 100 % $D_2O$ over the course of two days to allow bacteria to adapt to the deuterated medium. The main culture in M9 medium, which was prepared with 100 % $D_2O$, underwent the same procedure as described for (1). Isotope labeling in the final preculture and the main culture was achieved using $^{15}NH_4Cl$ and D-$^{13}C_6$-$^{2}H_7$-glucose.

For (3), the same protocol as for (2) was followed up until the main culture except for using D-$^{2}H_7$-glucose. The main culture was shaken until it reached an $OD_{600}$ of 0.6 at which point 100 mg $L^{-1}$ sodium [3,3-$^{2}H_2$]([3,5-$^{13}C_2$; 2,4,6-$^{2}H_3$] phenyl) pyruvate and 50 mg $L^{-1}$ sodium [3,3-$^{2}H_2$]([3,5-$^{13}C_2$; 2,6-$^{2}H_2$] 4-hydroxyphenyl) pyruvate (*2*) were added. After shaking for 1 h, expression was induced with 1 mM IPTG. The culture was shaken at 20 °C and 200 rpm overnight and harvested the next morning as described above.

The cell pellet was resuspended in 10 mL purification buffer (10 mM Tris, 1 mM ethylenediaminetetraacetic acid (EDTA), pH 7.5) and lysis was achieved by a heat shock at 80 °C for 10 min. After 5 min on ice, the suspension was centrifuged for 40 min at 4 °C and 47 850 × g. The supernatant was loaded onto a RESOURCE Q column (Cytiva) and eluted in purification buffer with a gradient of 0 to 1 M NaCl in 10 column volumes. The fractions containing GB1 were concentrated with an Amicon Ultra Centrifugal filter with 3 kDa molecular weight cutoff and size exclusion chromatography was performed on a HiLoad 16/600 Superdex 75 pg column (Cytiva) in crystallization buffer (50 mM sodium phosphate, pH 6.5).

##### Crystallization

Crystallization was achieved by dialysis (*3*) in a self-made dialysis button consisting of the upper part of a microtube and a dialysis membrane with 3.5 kDa molecular weight cutoff (SnakeSkin) (Fig. S1A). The protein in crystallization buffer was concentrated to 30 mg $mL^{-1}$ and 120 µL were loaded into the cap of the microtube. The dialysis button was closed with the dialysis membrane and placed into

10 mL of reservoir solution (50 % 2-methyl-2,4-pentanediol (MPD), 25 % isopropanol) (Fig. S1B). Dialysis was performed under constant agitation at 4 °C. Microcrystals started to grow after one to two days (Fig. S1C).

For NMR samples, the microcrystals were filled into a 1.3 mm or 1.9 mm (see Tab. S2) magic angle spinning (MAS) rotor (Bruker) by centrifugation (10 min at 4000 × g) after seven days.

To obtain crystals for X-ray diffraction (XRD), dialysis was performed for a total duration of two weeks, after which large rod-shaped crystals appeared.

For the success of crystallization, avoiding $Cl^-$ ions in the crystallization buffer is crucial (*4*). We also found that using an ultracentrifuge to transfer the crystals into the NMR rotor often leads to polymorphic samples. Although we did not investigate this systematically, we suspect that the vacuum in this kind of centrifuge leads to increased isopropanol evaporation, and we achieved more consistent results by using a tabletop centrifuge.

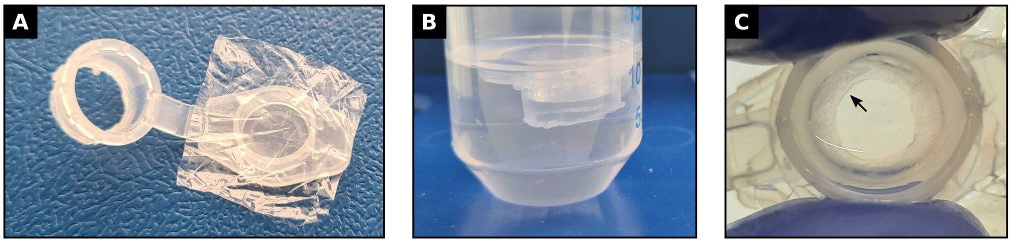

**Figure S1: A** Dialysis button made from the upper part of a microtube and dialysis membrane. **B** Dialysis button in reservoir solution. **C** Dialysis button with microcrystals forming a ring on the dialysis membrane.

**IgG:GB1 complex**

The sample of the IgG:GB1 complex was prepared as described previously (*5*). In short, lyophilized IgG from human serum (Sigma-Aldrich I4506) was dissolved in 50 mM sodium phosphate, pH 6.5 to a concentration of 0.15 mM. The antibody was mixed with 0.3 mM GB1 in the same buffer in a volume ratio of 1:1. The complex precipitates upon mixing and was filled into an MAS NMR rotor by centrifugation.

#### 1.2 Structure determination

Crystals were harvested at 4 °C, mounted on a MiTeGen cryoloop, and cooled to 100 K in liquid nitrogen without any additional cryoprotectant. Data collection was performed on the BM07-FIP2 beamline at the European Synchrotron Radiation Facility (ESRF) at 100 K. Two datasets, each comprising 1800 images, were recorded with 0.2° rotation and 0.1 s exposure per image. The images were collected at an energy of 12.657 keV from two different positions on the same crystal (see Fig. S2).

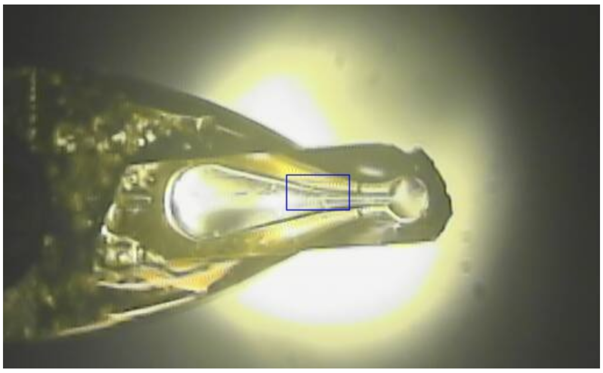

**Figure S2:** Crystal of GB1 (800x350x200 $\mu m^3$) mounted on a crystallization loop.

All diffraction frames of the two data sets were processed together using AUTOPROC (*6*), data were integrated in XDS (*7*), and the integrated intensities were scaled and merged in AIMLESS and POINTLESS, as implemented in CCP4 (*8*). AlphaFold2(*9*) was used to predict an initial model, which was used for molecular replacement with PHASER (*10*). The model was optimized through iterative rounds of model building in COOT (*11*) and refinement in PHENIX.REFINE (*12*). At all stages of the refinement, translation-libration-screw-rotation (TLS) was applied. Data collection and refinement statistics are reported in table S1. The structure and associated structure factor amplitudes were deposited in the Protein Data Bank (PDB) after validation with MolProbity (*13*) under the PDB code 9I2I.

**Table S1:** X-ray data collection and refinement statistics.

| **Data collection** | |
|---|---|
| Beamline | BM07-FIP2 (ESRF) |
| Space group | C2 |
| Cell dimensions | |
| a, b, c (Å) | 77.48, 35.18, 49.94 |
| $\alpha, \beta, \gamma$ (°) | 90.00 122.51 90.00 |
| Resolution (Å) | 42.11 – 1.08 (1.21 – 1.08)* |
| $R_{\mathrm{pim}}$ | 0.012 (0.259) |
| $I/\sigma I$ | 22.6 (2.0) |
| Completeness ellipsoidal (%) | 86.5 (45.3) |
| Completeness spherical (%) | 52.4 (9.3) |
| Redundancy | 13.5 (13.2) |
| **Refinement** | |
| Resolution (Å) | 20.82 – 1.08 (1.12 – 1.08)* |
| No. reflections | 25004 (92) |
| $R_{\mathrm{work}}/R_{\mathrm{free}}$ | 0.180 / 0.209 |
| No. atoms | |
| Protein | 902 |
| Ligand/ion | 1 |
| Water | 147 |
| B-factors | |
| Protein | 25.72 |
| Ligand/ion | 32.26 |
| Water | 41.11 |
| R.m.s. deviations | |
| Bond lengths (Å) | 0.006 |
| Bond angles (°) | 0.72 |
| Ramachandran | |
| Favored (%) | 98.08 |
| Allowed (%) | 1.92 |
| Clashscore | 1.68 |
| PDB ID | **9I2I** |

* Values in parentheses are for the highest-resolution shell.

#### 1.3 Small-angle X-ray scattering experiments

Samples for XRD were prepared as for the NMR experiments and placed inside borosilicate capillaries with 1 mm outer diameter, 0.01 mm wall thickness, and 80 mm length (Hilgenberg). The samples were measured on a XEUSS 3.0 HR (Xenocs SAS, France) in $< 0.1$ mbar vacuum using radiation from a microfocus Cu source collimated with a 3D multilayer mirror and shaped by scatterless slits, and an Eiger2

1M pixel array detector. Several sample-detector distances and collimation settings were tested for optimal data quality. A total of 36 frames of 120 s exposure each from multiple places in a capillary were cumulated to achieve good orientation statistics. The frames were cumulated and reduced in the XSACT suite (*14*). No background or buffer subtraction was done since all other scatterers (capillary, buffer) showed featureless patterns.

Analysis of the data was done using GSASII (*15*). We refined the unit cell parameters and the sample displacement following the approach of Le Bail for the peak intensities (*16*). An angle-independent Gaussian and a domain size-related Lorentzian term were used to fit the peak shapes and a polynomial background.

Refinement of the parameters resulted in $a$ = 77.9 Å, $b$ = 35.3 Å, $c$ = 51.4 Å, and $\beta$ = 122.3°, very close to the values obtained by single-crystal XRD (see table S1).

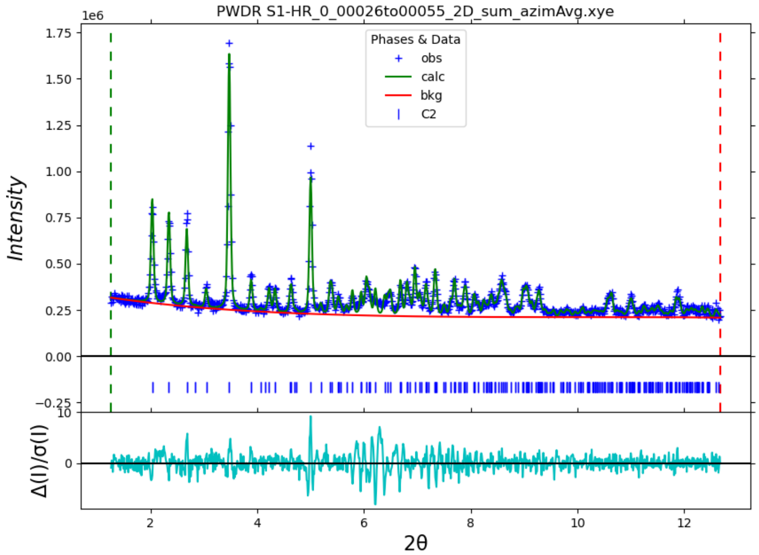

**Figure S3:** Powder X-ray diffraction of GB1$_{QDD}$ microcrystals.

#### 1.4 NMR spectroscopy

MAS NMR experiments were recorded on a Bruker Avance Neo spectrometer operating at 16.44 T (700 MHz $^{1}$H Larmor frequency) with triple channel ($^{1}$H, $^{13}$C, $^{15}$N) probe heads for 1.3 mm or 1.9 mm MAS rotors. All measurements were performed with $^{1}$H detection and cross-polarization (CP) based transfers. The following sequences used in this work are implemented in the ssNMRlib library (*17*): 2D hNH, 3D hCONH, 3D hCOcaNH, 3D hCANH, 3D hCAcoNH, 3D hcaCBcaNH, 3D hcaCBcacoNH, 3D HhNH RFDR, pseudo-3D hNH $R_1$ and pseudo-3D hNH $R_{1\rho}$. Experiments used to study the aromatic $^{1}$H–$^{13}$C signals of specifically (CH)$^{\epsilon}$ labeled Phe and Tyr residues are shown in Figure S4 (not currently implemented in NMRlib). They all contain a $^{13}$C EBURP2 pulse (*18*) to selectively excite the aromatic C$^{\epsilon}$ signal and suppress the natural-abundance carbon background that results particularly from cross-polarization from amide hydrogens (which spectrally overlap with those of aromatic hydrogens) and C$^{\alpha}$ or C' carbons. Additionally, the water suppression (ws) element was moved in front of the $^{13}$C chemical-shift evolution period $t_1$ to avoid cross-peaks between C$^{\epsilon 1}$ and C$^{\epsilon 2}$ that may arise from exchange dynamics or magnetization transfer (proton-driven spin diffusion). In all cases, the water-suppression element consisted of a composite-pulse decoupling sequence with constant amplitude and a pseudo-random variation of the pulse durations and phases. The determination of the sample temperature was performed on an external sample (1.3 or 1.9 mm rotor) containing lyophilized protein rehydrated with a solution of 25 mM TmDOTP and 10 mM 2,2-dimethyl-2-silapentane-5-sulfonate sodium salt (DSS) in 80 % $D_2O$ for each measurement condition (*19*). The experimental conditions are summarized in Table S2.

**Assignment experiments**

Six 3D spectra (hCONH, hCOcaNH, hCANH, hCAcoNH, hcaCBcaNH, hcaCBcacoNH) were recorded to assign the resonances of $H^N$, $N^H$, C', $C^\alpha$ and $C^\beta$ atoms of GB1 in the crystal.
A 3D HhNH experiment with $^1H-^1H$ radiofrequency-driven recoupling (RFDR) was used to assign aromatic ($H^\epsilon$) protons of $(CH)^\epsilon$-labeled Phe and Tyr by probing their proximity to amide ($H^N$) protons.
A 2D HcH Exchange spectroscopy (EXSY) experiment was recorded to probe potential exchange between $^{13}C^\epsilon$ atoms in tyrosines in the IgG:GB1 complex (Fig. S4B). The exchange delay $\tau$ of 256 ms was applied while magnetization was stored $^{13}C$ nuclei after the indirect $^1H$ evolution period.

**Experiments to probe dynamics**

Experiments to probe amide $^{15}N$ and aromatic $^{13}C$ longitudinal $R_1$ and rotating frame relaxation $R_{1\rho}$ were performed as pseudo-3D spectra. The pseudo dimension was used to vary the relaxation delay $\tau$ or the duration of the spinlock pulse (SL) for $R_1$ and $R_{1\rho}$, respectively (Fig. S4C,D). The delays and pulse lengths used can be found in Figures S11, S16, S17, and S18.
EXSY experiments were performed to study the timescale of tyrosine ring flips for which separate signals for $H^{\epsilon 1}$-$C^{\epsilon 1}$ (site a) and $H^{\epsilon 2}$-$C^{\epsilon 2}$ (site b) could be observed (Fig. S4E). If a ring flips during the longitudinal $C^z$ mixing time $\tau$, sites a and b exchange, resulting in the cross peaks $H^{\epsilon 2}$-$C^{\epsilon 1}$ (ab) and $H^{\epsilon 1}$-$C^{\epsilon 2}$ (ba) in the final spectrum. The experiment was implemented as a pseudo-4D experiment with two frequency dimensions ($^1H$ and $^{13}C$), and two dimensions related to the EXSY: along one dimension, a variable exchange delay $\tau$ allowed monitoring the build-up (due to the exchange process) and subsequent decay (due to $R_1$ relaxation, i.e. incoherent processes, and possibly spin diffusion, i.e. coherent transfer of magnetization to other nuclei) of cross-peak intensities. In the fourth dimension, the position of the exchange delay $\tau$ was alternated, to be either before or after the $^{13}C$ chemical-shift evolution period $t_1$. In the former case, only diagonal peaks are observed, and their intensity decays over $\tau$. In the latter case, the build-up of exchange cross-peaks is observed, and the intensities of diagonal and cross-peaks are subject to the same decay as in the reference experiment. When subtracting the two spectra, diagonal peaks are eliminated, facilitating the quantification of cross-peaks. As part of the exchange delay, an additional water suppression element with a constant duration was implemented, making the minimum possible $\tau$ 10 ms. We verified whether the observed cross peaks may result from spin diffusion rather than from ring flips, by repeating the experiments at different MAS frequencies $\nu_r$ (30 kHz and 39 kHz) while keeping the sample temperature constant. Spin diffusion, but not ring-flip dynamics, are expected to be MAS frequency dependent.
Rotational-Echo Double Resonance (REDOR) experiments were performed with a shift $\Delta$ of the first inversion pulse in each rotor period $\tau_r$ to scale down the dipolar coupling and allow better sampling (Fig. S4F) (*20, 21*). The dephasing curve was sampled at time points $2\tau_r + (n-1)2\tau_r$ with $n = 1-15$. The incrementation of rotor periods was implemented as a third dimension. The REDOR block was followed by a short z-filter to suppress unwanted coherences that might have built up during the recoupling period. The reference experiment in which the inversion pulses on the $^1H$ channel were replaced with equally long delays was sampled at $n = 1, 7, 13$, and the missing reference intensities were obtained from a linear fit of these points. This approach allows spending more time on collecting the more informative data points of the recoupling experiment because the decay in such highly deuterated samples is slow and sensitivity high, such that the fit of the data is very robust.

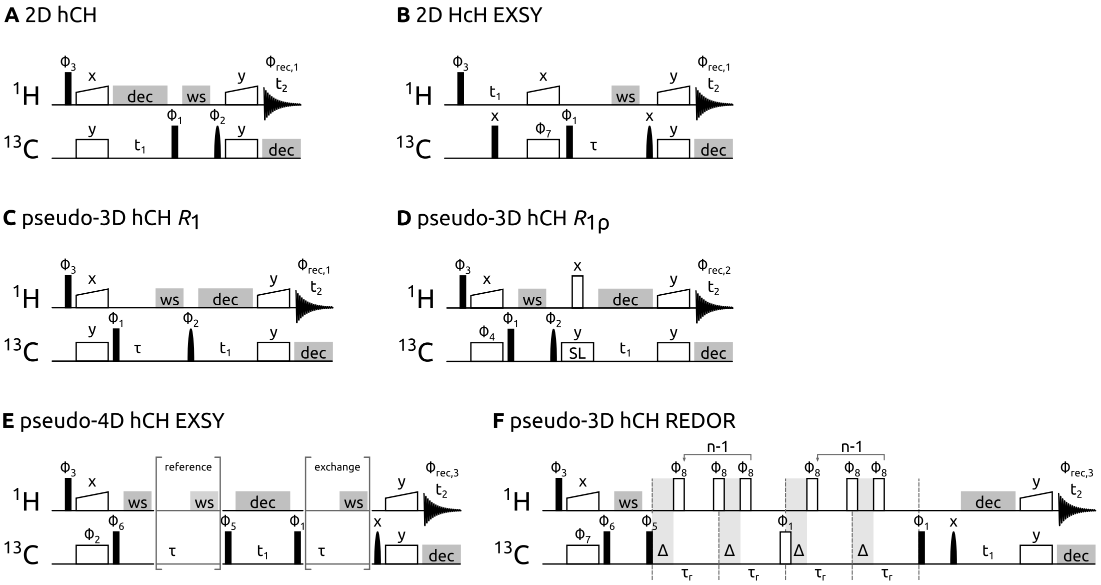

**Figure S4:** Pulse sequences used in this study for aromatics. Narrow black rectangles denote 90° pulses, open rectangles denote 180° pulses and rounded black symbols denote selective $^{13}$C excitation pulses (EBURP2). Grey rectangles indicate decoupling (dec) and water suppression (ws) elements. Wide open symbols denote cross-polarization elements. Indirect and direct acquisition times are indicated with $t_1$ and $t_2$ respectively. Phase cycles are: $\Phi_1$ = x,-x; $\Phi_2$ = 2(x),2(-x); $\Phi_3$ = 4(y),4(-y); $\Phi_4$ = 8(y),8(-y); $\Phi_5$ = 8(x),8(-x); $\Phi_6$ = 16(x),16(-x); $\Phi_7$ = 2(y),2(-y); $\Phi_8$ = XY-8; $\Phi_{rec,1}$ = y,-y,-y,y, -y,y,y,-y; $\Phi_{rec,2}$ = y,-y,-y,y, 2(-y,y,y,-y), y,-y,-y,y; $\Phi_{rec,3}$ = y,-y,-y,y, 2(-y,y,y,-y), y,-y,-y,y, -y,y,y,-y, 2(y,-y,-y,y), -y,y,y,-y.

**Table S2:** Experimental NMR parameters.

| Figures | Experiment | Fig. S4 | Sample | Probe [mm] | $\nu_r$ [kHz] | $T_{set}$ [K] | $T_{sample}$ [K] | $t_{exp}$ [h] | $t_1/t_2/t_{dir}$[a][ms] | w.s.[b][kHz] | dec. $^1H/^{13}C/^{15}N$[b][kHz] | Comment |
|---|---|---|---|---|---|---|---|---|---|---|---|---|
| 2A | hNH | | (2), crystal | 1.9 | 40 | 245 | 305 | 0.5 | 32/-/15 | 16 | 12.5/-/5 | |
| S5 | hCANH | | (2), crystal | 1.9 | 40 | 245 | 305 | 9 | 6/22/15 | 16 | 12.5/-/5 | NUS: 35 %[d] |
| S5 | hCAcoNH | | (2), crystal | 1.9 | 40 | 245 | 305 | 16 | 6/22/15 | 16 | 12.5/-/5 | NUS: 60 %[d] |
| S5 | hcaCBcaNH | | (2), crystal | 1.9 | 40 | 245 | 305 | 23.5 | 4.5/22/15 | 16 | 12.5/-/5 | NUS: 70 %[d] |
| S5 | hcaCBcacoNH | | (2), crystal | 1.9 | 40 | 245 | 305 | 47 | 4.5/22/15 | 16 | 12.5/-/5 | NUS: 70 %[d] |
| S5 | hCONH | | (2), crystal | 1.9 | 40 | 245 | 305 | 12 | 24/22/15 | 18 | 12.5/-/5 | NUS: 34 %[d] |
| S5 | hCOcaNH | | (2), crystal | 1.9 | 40 | 245 | 305 | 21.5 | 24/22/15 | 16 | 12.5/-/5 | NUS: 60 %[d] |
| 2C, S11 | hNH $R_1$ | | (3), crystal | 1.9 | 39 | 245 | 304 | 60.5 | 32/-/15 | 16 | 100/-/5 | |
| 2C, S11 | hNH $R_{1\rho}$ | | (3), crystal | 1.9 | 39 | 245 | 304 | 12.5 - 49 | 32/-/15 | 16 | 100/-/5 | $\nu_{SL}$ = 15, 25, 27.5, 30, 32.5, 35 kHz |
| 2D | hCH | | solution | | | 288 | 288 | | 28/-/61 | | | |
| 2D | hCH | A | (3), crystal | 1.3 | 39 | 225 | 288 | 0.5 | 8/-/15 | 25 | 12.5/5/- | |
| 2D, S10 | hCH | A | (3), IgG:GB1 | 1.9 | 30 | 245 | | 73.5 | 9/-/15 | 18.5 | 12.5/15/- | |
| S7 | HNH RFDR | | (3), crystal | 1.9 | 40 | 250 | 307 | 45.5 | 22/4/15 | 16 | 12.5/-/5 | |
| S10 | HcH EXSY | B | (3), IgG:GB1 | 1.9 | 39 | 245 | 304 | 16.5 | 4.5/-/30 | 16 | 7.5/5/- | |
| 3A, S13B | EXSY | E | (3), crystal | 1.9 | 30 | 271 | 304 | 50 | 10/-/15 | 16/18.5 | 100/5/- | |
| 3A | EXSY | E | (3), crystal | 1.9 | 39 | 245 | 304 | 61.5 | 10/-/15 | 16/18.5 | 100/5/- | |
| 3A, S13A | EXSY | E | (3), crystal | 1.3 | 39 | 225 | 288 | 133 | 8/-/15 | 16/18.5 | 100/5/- | |
| 4, S14 | hCH REDOR | F | (3), crystal | 1.3 | 39 | 256 | 304 | 138 | 8/-/15 | 16 | 12.5/5/- | $\Delta$ = 3 µs |
| 4, S15 | hCH REDOR | F | (3), IgG:GB1 | 1.9 | 39 | 245 | 304 | 92.5 | 4/-/30 | 16 | 7.5/5/- | $\Delta$ = 2.5 µs |
| 4, S17, S19 | hCH $R_{1\rho}$ | D | (3), crystal | 1.9 | 39 | 245 | 304 | 13.5 - 26.5 | 10/-/15 | 16 | 100/5/- | $\nu_{SL}$ = 15, 25, 27.5, 30, 32.5, 35 kHz |
| 4, S16, S19 | hCH $R_1$ | C | (3), crystal | 1.3 | 39 | 225 | 288 | 48.5 | 8/-/15 | 16 | 100/5/- | |
| 4, S16, S19 | hCH $R_{1\rho}$ | D | (3), crystal | 1.3 | 39 | 225 | 288 | 21.5 - 26.5 | 8/-/15 | 16 | 100/5/- | $\nu_{SL}$ = 15, 25, 27.5, 30, 32.5, 35 kHz |
| 4, S18, S19 | hCH $R_1$ | C | (3), IgG:GB1 | 1.9 | 39 | 245 | 304 | 50.5 | 30/-/- | 16 | 12.5/5/- | |
| 4, S18, S19 | hCH $R_{1\rho}$ | D | (3), IgG:GB1 | 1.9 | 39 | 245 | 304 | 15.5 | 30/-/- | 16 | 12.5/5/- | $\nu_{SL}$ = 15, 25, 27.5, 30, 32.5, 35 kHz |

[a] Acquisition time in indirect ($t_1/t_2$) and direct ($t_{dir}$) dimension. [b] Water suppression achieved with a composite pulse decoupling scheme based on a modified MISSISSIPPI sequence (*22*). The second value for longitudinal exchange experiments corresponds to the ws block during the mixing time. [c] Composite pulse decoupling during acquisition with swfTPPM (*23*) for $^1H$ and WALTZ-16 (*24*) for $^{13}C$ and $^{15}N$.
[d] Amount of non-uniform sampling.

#### 1.5 NMR Data Analysis

All spectra were processed with Bruker Topspin 4.1.4 and then converted to UCSF format with the bruk2ucsf program provided in Sparky (*25*). The analysis of the experiments to investigate dynamics was done with scripts using the Nmrglue package for Python (*26*). All spectra were indirectly referenced to DSS via MPD (*27*).

##### Backbone assignment

Non-uniform sampling (NUS) reconstruction and processing of backbone assignment spectra was performed in Topspin before conversion to UCSF format. Peaks were picked in CcpNmr Analysis Version 3 (*28*) and exported for automated resonance assignment by FLYA (CYANA v3.98.15) (*29*). Chemical-shift tolerances were set to 0.1 ppm for $^{1}$H and 0.4 ppm for $^{13}$C and $^{15}$N resonances. Minor manual corrections to the automated assignment were made in CcpNmr. Representative sections of the 3D spectra are shown in Figure S5. The completeness of the assignment is visualized in Figure S6 and all assigned chemical shifts can be found in Table S3.

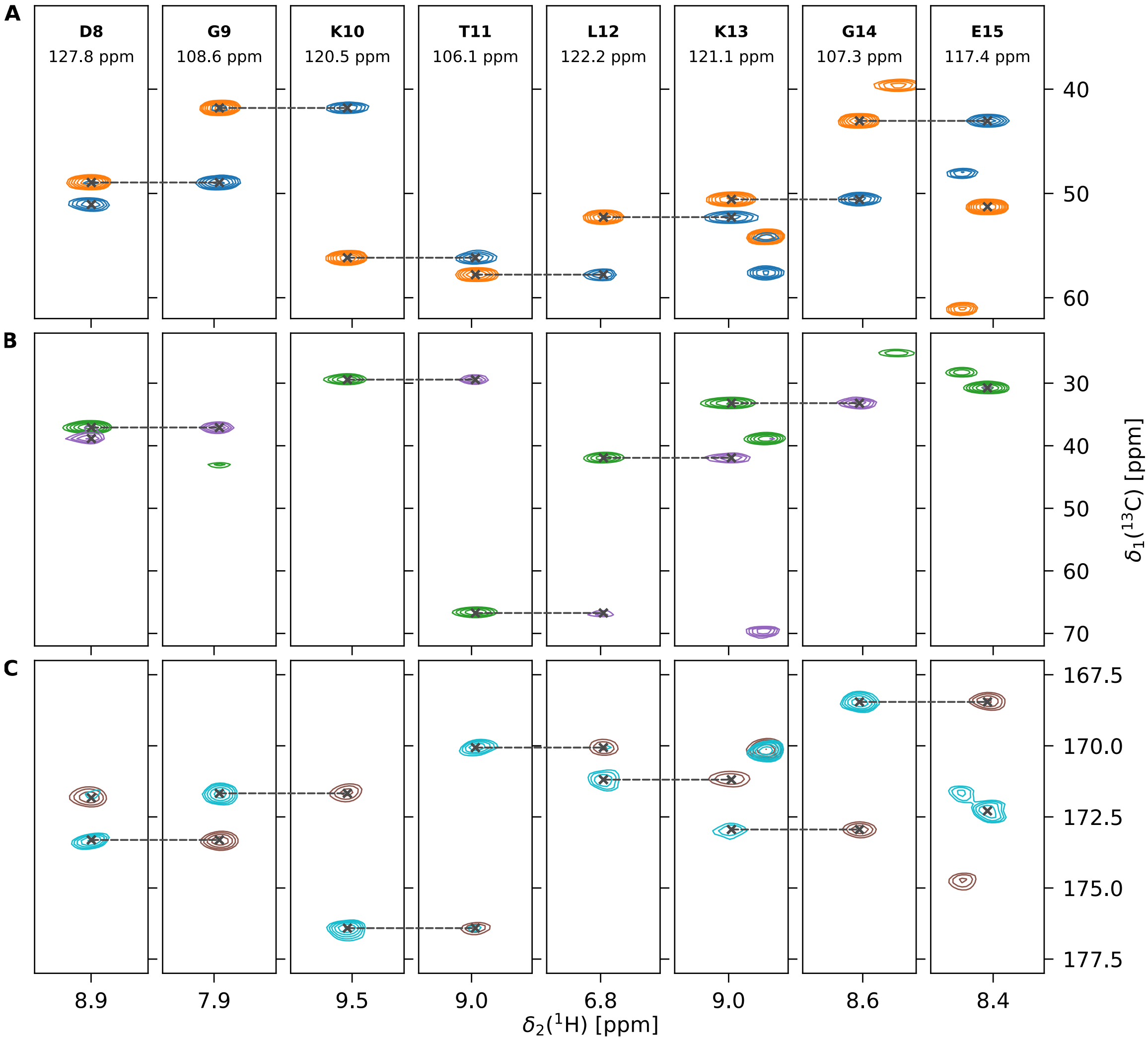

**Figure S5:** Representative sections of spectra for backbone assignment for residues D8 to E15. Each column shows the plane at the assigned $^{15}$N frequency of one residue as indicated at the top. **A** hCANH (orange) and hCAcoNH (blue). **B** hcaCBcaNH (green) and hcaCBcacoNH (purple). **C** hCOcaNH (cyan) and hCONH (brown).

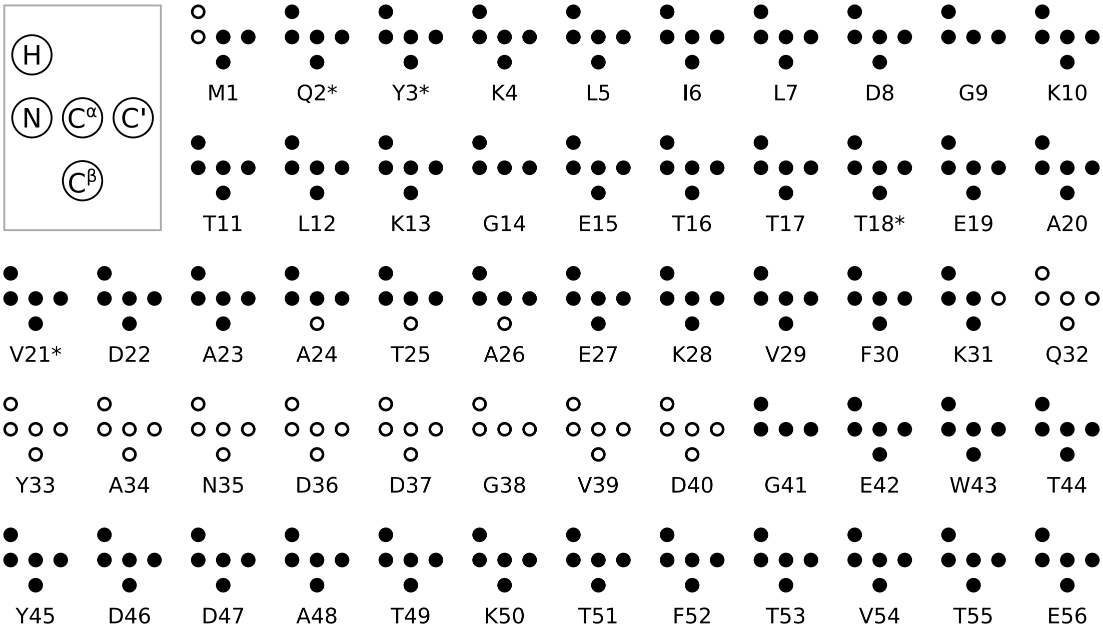

**Figure S6:** Completeness of the backbone assignment for each residue. Empty and filled circles indicate unassigned and assigned atoms, respectively. Residues that have two sets of resonances are marked with an asterisk.

**Table S3:** Chemical shift assignments in [ppm]. Residues with two rows have two sets of resonances.

| Residue | $H^N$ | $N^H$ | C' | $C^\alpha$ | $C^\beta$ | Residue | $H^N$ | $N^H$ | C' | $C^\alpha$ | $C^\beta$ |
|---|---|---|---|---|---|---|---|---|---|---|---|
| M1 | | | 168.04 | 51.53 | 28.85 | E27 | 8.49 | 116.23 | 174.62 | 56.55 | 25.37 |
| Q2 | 8.10 | 123.38 | 171.56 | 52.46 | 26.57 | K28 | 6.88 | 116.05 | 176.67 | 56.42 | 28.52 |
| | 8.31 | 125.58 | 171.76 | 53.03 | 26.87 | V29 | 7.50 | 119.54 | 176.01 | 63.13 | 28.13 |
| Y3 | 8.65 | 119.83 | 172.16 | 54.15 | 40.65 | F30 | 8.64 | 118.38 | 175.89 | 54.51 | 34.60 |
| | 8.87 | 123.60 | 172.28 | 54.15 | 39.92 | K31 | 8.85 | 120.23 | | 56.82 | 28.52 |
| K4 | 9.20 | 122.33 | 170.19 | 51.90 | 32.09 | Q32 | | | | | |
| L5 | 8.73 | 126.84 | 171.84 | 50.00 | 39.45 | Y33 | | | | | |
| I6 | 9.28 | 125.89 | 172.03 | 56.83 | 34.52 | A34 | | | | | |
| L7 | 8.95 | 126.40 | 171.83 | 51.06 | 38.85 | N35 | | | | | |
| D8 | 8.90 | 127.76 | 173.31 | 48.95 | 37.08 | D36 | | | | | |
| G9 | 7.86 | 108.58 | 171.67 | 41.80 | | D37 | | | | | |
| K10 | 9.53 | 120.55 | 176.41 | 56.16 | 29.43 | G38 | | | | | |
| T11 | 8.97 | 106.12 | 170.06 | 57.79 | 66.73 | V39 | | | | | |
| L12 | 6.78 | 122.23 | 171.19 | 52.27 | 41.93 | D40 | | | | | |
| K13 | 8.98 | 121.10 | 172.94 | 50.57 | 33.19 | G41 | 7.60 | 104.34 | 169.27 | 41.64 | |
| G14 | 8.62 | 107.31 | 168.46 | 43.04 | | E42 | 8.39 | 119.17 | 173.60 | 52.29 | 27.90 |
| E15 | 8.44 | 117.38 | 172.29 | 51.28 | 30.75 | W43 | 8.85 | 131.80 | 173.63 | 54.96 | 26.06 |
| T16 | 8.89 | 115.48 | 169.20 | 57.63 | 66.56 | T44 | 9.56 | 114.84 | 170.14 | 57.61 | 69.49 |
| T17 | 8.20 | 111.82 | 171.49 | 57.34 | 70.14 | Y45 | 8.73 | 121.29 | 170.19 | 54.19 | 38.87 |
| T18 | 8.99 | 111.77 | 168.84 | 59.49 | 66.74 | D46 | 7.78 | 127.60 | 172.22 | 48.85 | 40.42 |
| | 9.09 | 113.33 | 168.56 | 59.23 | 66.77 | D47 | 8.89 | 125.22 | 175.73 | 53.90 | 39.93 |
| E19 | 7.78 | 124.68 | 172.75 | 51.47 | 28.40 | A48 | 8.20 | 118.17 | 176.44 | 51.82 | 14.98 |
| A20 | 9.51 | 126.15 | 174.89 | 47.98 | 19.84 | T49 | 6.90 | 101.86 | 172.24 | 57.22 | 67.25 |
| V21 | 8.61 | 117.14 | 171.62 | 60.77 | 28.31 | K50 | 7.83 | 123.25 | 172.29 | 52.92 | 26.02 |
| | 9.17 | 114.73 | 172.43 | 60.32 | 28.43 | T51 | 7.57 | 110.97 | 172.21 | 59.26 | 69.21 |
| D22 | 7.24 | 114.27 | 173.01 | 49.87 | 38.85 | F52 | 10.38 | 130.16 | 171.93 | 54.23 | 38.99 |
| A23 | 9.54 | 122.14 | 176.51 | 51.40 | 14.07 | T53 | 9.08 | 117.94 | 169.99 | 58.84 | 67.73 |
| A24 | 7.89 | 120.30 | 178.40 | 51.55 | | V54 | 8.33 | 124.27 | 170.83 | 55.79 | 28.70 |
| T25 | 8.29 | 116.73 | 172.91 | 64.42 | | T55 | 8.11 | 122.94 | 171.48 | 58.52 | 67.59 |
| A26 | 7.46 | 123.36 | 174.56 | 51.94 | | E56 | 7.97 | 133.24 | 177.45 | 54.49 | 29.62 |

#### Assignment of tyrosine side chains in GB1 crystals

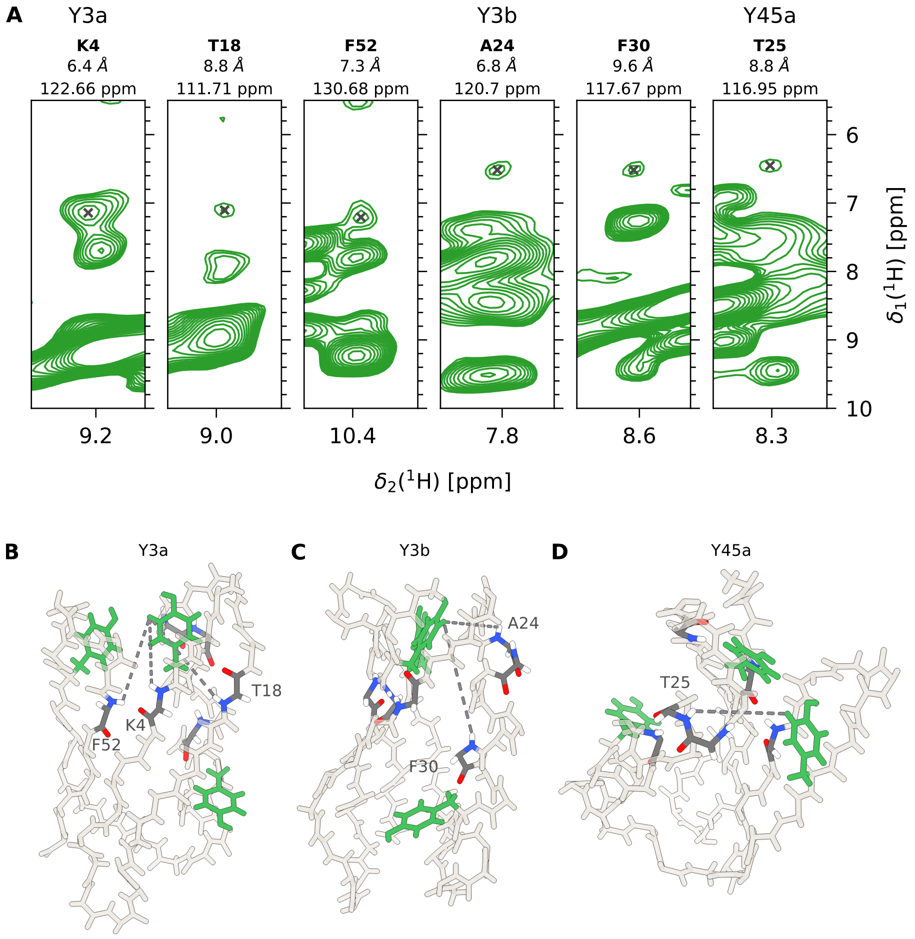

**Figure S7:** Assignment of $(CH)^{\epsilon}$-labeled tyrosines in $GB1_{QDD}$ crystals by a 3D HNH RFDR spectrum. (A) Sections of a 3D HNH RFDR spectrum showing cross peaks between the $^{1}H$ frequency of $(CH)^{\epsilon}$-labeled tyrosines and the $^{1}H$ frequency of backbone amide proton signals. The header of each strip indicates the assigned tyrosine signal, the residue of the backbone signal, the distance between the two atoms, and the $^{15}N$ frequency of the shown plane. (B-D) 3D representation of $GB1_{QDD}$ indicating the observed cross peaks from (A) as connections between the atoms. The backbone is represented by beige, and the three tyrosines by green sticks. Residues observed in cross peaks are highlighted in dark gray. Each panel shows the connections of one tyrosine as indicated on top.

#### Determination of ring flip rates from EXSY experiments

The pseudo-4D spectrum was processed as individual 2D planes resulting in a reference and an exchange spectrum for each exchange time. Peak intensities were obtained by fitting a two-dimensional Gaussian lineshape with functions provided in the Nmrglue package. Intensities of the diagonal and cross peaks were extracted in the reference and difference spectrum respectively. For residue Y45, the four signals (A, B, AB, BA) were fitted as described in Farrow *et al.*, 1994 (*30*) to obtain the rate of the ring flip $k_{\text{flip}}$. The longitudinal relaxation rates $R_{1,\text{A}}$ and $R_{1,\text{B}}$ were set to the determined values from the relaxation experiment and the populations were set to $p_{\text{A}} = p_{\text{B}} = 0.5$. Due to significant signal overlap of Y3 and Y33, we extracted $k_{\text{flip}}$ of residue Y3 only from the build-up of the cross-peak BA intensities $I$ according to $I(t) = I(0)(1 - \exp[-2k_{\text{flip}}t]) \exp[-R_{1,\text{B}}t]$. The fitted curves are shown in Figure S13. Errors were determined by Monte Carlo analysis. 500 noisy data sets were generated using the spectral noise level defined as the standard deviation of intensities across a spectral region without any signal. The error of each fit parameter was estimated as the standard deviation of that given parameter across the fits of these synthetic data sets.

**Determination of relaxation rate constants**

Peak intensities were extracted from a series of 2D planes with different relaxation delays ($R_1$) or spin-lock pulse length ($R_{1\rho}$) as described above. Relaxation rate constants were obtained by an exponential fit of the intensities. Error analysis was performed as described above. All fits are shown in Figures S16, S17, S18.

**Detectors**

Relaxation rates were analyzed using the Detectors approach as implemented in the package Detectorist (*31*). Sensitivities were calculated for each of the relaxation rates measured. For the site-selective $^{13}C-^{1}H$ Tyrs, the sensitivity was calculated assuming a $^{13}C-^{1}H$ dipolar coupling at a distance of 1.09 Å, and a CSA tensor with principal components 180, 143, and 27 ppm. For the backbone $^{15}N$ sites, the sensitivity was calculated assuming the directly bonded $^{1}H$ at a distance of 1.02 Å, and principal components of the CSA tensor of 229.2, 78.6, 52.3 ppm. Each sensitivity was digitized using 200 time points, logarithmically spaced between 100 fs/rad and 1 ms/rad. It should be noted that owing to the form of the spectral density, the correlation times involved are naturally in units of seconds per radian. For comparison with results from EXSY, these have been explicitly converted into the base unit of seconds.
Detector optimization was performed using the Singular Value Decomposition method (*32*). Specifically, a matrix, $\boldsymbol{M}$, was produced whereby each of the $N$ rows represents the sensitivity of a corresponding relaxation rate. The singular value decomposition of this matrix was taken:

$$\boldsymbol{M} = \boldsymbol{U}\boldsymbol{\Sigma}\boldsymbol{V}'. \tag{1}$$

The singular values (diagonal entries in $\boldsymbol{\Sigma}$) were sorted, the matrices truncated to include $k$ singular values ($k$ was varied from 2 to 7 for model selection). Linear programming was performed to optimize linear combinations of the orthogonal singular vectors in the truncated $\boldsymbol{V}'_k$ to identify well-formed detectors. The coefficients of $\boldsymbol{V}'_k$ giving rise to detectors then form the rows of the transformation matrix $\boldsymbol{Q}$, which can be used to determine the matrix $\boldsymbol{r}$, containing 'detector vectors':

$$\boldsymbol{r} = \boldsymbol{U}_k\boldsymbol{\Sigma}_k\boldsymbol{Q}^{-1}, \tag{2}$$

for which each of the $N$ columns, $\boldsymbol{r}_i$, gives a linear combination of $\leq k$ detectors giving the corresponding relaxation rate. The detector responses may then be calculated using a Non-Negative Least Squares to solve for the detector responses, $\boldsymbol{\rho}$:

$$\boldsymbol{\rho} = \begin{bmatrix} \boldsymbol{r}_1/e_1 \\ \vdots \\ \boldsymbol{r}_N/e_N \end{bmatrix}^{-1} \frac{\boldsymbol{R}}{\boldsymbol{e}}, \tag{3}$$

where $\boldsymbol{R}$ and $\boldsymbol{e}$ are vectors containing the experimental relaxation rates and uncertainties, respectively. The detector sensitivities are obtainable directly from the product $\boldsymbol{Q}\boldsymbol{V}'_k$. The number of singular values to be included in the analysis was identified by performing the analysis for $k = 2$ to 7, back-calculating the relaxation rates, then choosing the number of singular values that gave the lowest median reduced chisq value.

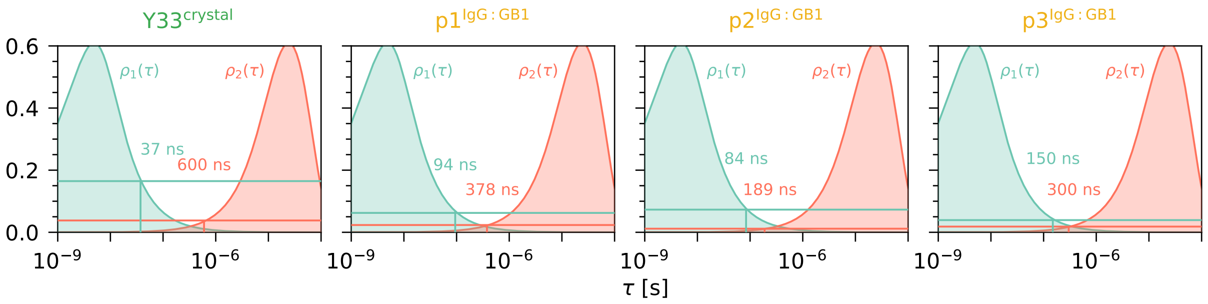

**Figure S8: Estimation of timescales for tyrosines Y33$^{\text{crystal}}$, and p1, p2, and p3 of the complex.** If a dynamic process has a correlation time $\tau$ equal to the maximum of the detector sensitivity, the response to this sensitivity equals $(1 - S^2)$, which is 0.609 for a ring flip. All responses we obtained are much lower than this value, indicating that none of the residues flip at a timescale corresponding to the maximum of one of the sensitivities. We know from the dipolar order parameters that Y33$^{\text{crystal}}$, p1, p2, and p3 flip faster than hundreds of microseconds. As ring flips faster than tens of nanoseconds are unlikely, we assume that these residues flip on a timescale that lies in the 'blind spot' between $\rho_1$ and $\rho_2$. This blind spot is due to a lack of experiments that can accurately probe dynamics on this timescale. Here, we show a close-up of $\rho_1$ (blue) and $\rho_2$ (red), which are scaled by $(1 - S^2) = 0.609$. The horizontal lines are the detector responses for the two sensitivities, respectively. The intercepts of the response and the sensitivity are the minimum and maximum timescales that define the range in which the ring should flip. Figure 4D of the main text shows this in a compacted way.

###### Determination of dipolar order parameters from REDOR experiments

Peak intensities were extracted as described above. The reference curve was linearly interpolated to obtain intensities $S_0$ at the same time points as in the dephasing experiment $S_{\text{rec}}$. The REDOR curve was calculated as $\Delta S/S_0 = (S_0 - S_{\text{rec}})/S_0$. The observed dipolar coupling strength $\delta_D^{\text{obs}}$ and tensor asymmetry $\eta$ were determined by a two-dimensional grid search against simulated curves. The simulated curves with varying $\delta_D$ (0-50 000 Hz with a step size of 50 Hz) and $\eta$ (0-1 with a step size of 0.05) were generated using the GAMMA simulation package for numerical spin dynamics simulations (*33*). Chi-square minimization between the simulated and experimental curves was used to find the best-fit values for $\delta$ and $\eta$. Error determination was performed as described above. Dipolar order parameters $S$ were calculated with $S = (\delta_D^{\text{obs}}/\delta_D^{\text{rigid}})$ with $\delta_D^{\text{rigid}} = 46\,656$ Hz, corresponding to a $^1$H–$^{13}$C distance of 1.09 Å. The fits are shown in Figures S14 and S15.

#### 1.6 Computational details

###### Computational assays

Two distinct computational assays were constructed for GB1 in its crystal form and in an aqueous solution, using the visualization package VMD 1.9.4 (*34*), and the CHARMM-GUI utility (*35*). The crystal assay consisted of eight proteins and 2456 water molecules. The aqueous solution consisted of a single protein immersed in a bath of 6384 water molecules. To ensure electric neutrality of each assay, and mimic physiological conditions, NaCl was added at a concentration of 150 mM. After suitable thermalization, the dimensions of computational assays were approximately $a$ = 80.2 Å, $b$ = 36.4 Å, $c$ = 53.2 Å, with $\alpha$ = 90.0°, $\beta$ = 120.7°, $\gamma$ = 90.0°, for the crystal form, and $a$ = 58.7 Å, $b$ = 58.7 Å, $c$ = 58.7 Å, for the aqueous solution, corresponding to 16704 and 26432 atoms, respectively.

###### Molecular Dynamics Simulations

All molecular dynamics simulations were carried out using NAMD 3.0 (*36*), with the AMBER ff19SB force field for proteins and the OPC water model (*37*). Periodic boundary conditions were applied for all computational systems. The temperature was maintained at 298 K by means of a stochastic velocity rescaling thermostat (*38*) with a time parameter of 1 ps$^{-1}$ for coupling, and pressure at 1 bar, using the Langevin piston algorithm (*39*). All covalent bonds between heavy and hydrogen atoms were constrained to their equilibrium length with the Rattle algorithm (*40*). Water molecules were constrained to their equilibrium geometry with the Settle algorithm (*41*). Long-range electrostatic forces were computed with the particle-mesh Ewald algorithm (*42*), and a grid spacing of 1.2 Å. Short-range van der Waals and electrostatic interactions were smoothly truncated with a 10 Å spherical cut-off. Hydrogen-mass repartitioning was applied to the protein and surrounding lipids to allow longer integration time steps to be utilized (*43*). The Verlet-I/r-RESPA multiple time-stepping algorithm was employed to integrate the equations of motion with an effective time step of 4 fs for short-range interactions, and 8 fs for long-range interactions (*44*).

###### Free-Energy calculations

As a preamble to the free-energy calculations, the two computational assays underwent a suitable energy minimization, followed by 400 ns of thermalization bereft of geometrical restraints. The potentials of mean force (PMFs), $\Delta G(\chi_2)$, underlying the isomerization of the side chain of residues Y3, Y33 and Y45 about the $\chi_2$ torsional angle, in the crystal form of the protein and in the aqueous solution, were determined with the well-tempered metadynamics extended adaptive biasing force (WTM-eABF) algorithm (*45*, *46*). In a nutshell, it relies upon the integration of the average force acting on $\chi_2$, obtained from unconstrained molecular dynamics simulations. In the course of the simulation, a biasing force is estimated such that, once applied to the system, it yields a Hamiltonian devoid of an average force exerted along $\chi_2$. As a result, all values of $\chi_2$ are sampled with an equal probability, which, in turn, greatly improves the accuracy of the calculated free energies. The transition pathway spanned 360°, i.e., $-180° \leq \chi_2 \leq +180°$, and was discretized in bins 5° wide, where samples of the local force acting along $\chi_2$ were accrued. To mitigate the risk of deleterious nonequilibrium effects, no time-dependent bias was applied until a threshold of 10000 samples was reached (*47*). In consideration of the significant isomerization timescales measured experimentally, likely to be related to slowly relaxing degrees of freedom coupled to $\chi_2$, likely to hamper convergence of the free-energy calculation, a multiple-walker strategy was employed to improve ergodic sampling (*48*), with up to four walkers. The statistical error associated to the PMFs was estimated from the variance of the free-energy differences measured by each walker. Analysis of the trajectories was performed using VMD 1.9.4 (*36*).

#### 1.7 Considerations about dependency of rate constants on $\Delta H^\ddagger$ and $\Delta S^\ddagger$

In the main text we argued that the flatter dependency of the rate constants as a function of the inverse temperature, $\ln(k_{\text{flip}}) = f(1/T)$, in the crystal compared to solution (Fig. 5B) suggests that $\Delta H^{\ddagger,\text{solution}} \geq \Delta H^{\ddagger,\text{crystal}}$.

Here, we show that this conclusion follows simply from the Eyring equation, which is given by:

$$k = \frac{k_B T}{h} \exp(\frac{\Delta S^\ddagger}{R}) \exp(\frac{-\Delta H^\ddagger}{RT}) \tag{4}$$

which can be rearranged as follows:

$$\begin{aligned}\ln k &= \ln\left(\frac{k_B T}{h}\right) + \frac{\Delta S^\ddagger}{R} - \frac{\Delta H^\ddagger}{RT} \\ &= \ln k_B + \ln T - \ln h + \frac{\Delta S^\ddagger}{R} - \frac{\Delta H^\ddagger}{RT} \\ &= \left(\ln k_B - \ln h + \frac{\Delta S^\ddagger}{R}\right) + \ln T - \frac{\Delta H^\ddagger}{RT} \\ &= \left(\ln k_B - \ln h + \frac{\Delta S^\ddagger}{R}\right) + \ln T - \frac{\Delta H^\ddagger}{R} \cdot \frac{1}{T}\end{aligned} \tag{5}$$

With $c = \left(\ln k_B - \ln h + \frac{\Delta S^\ddagger}{R}\right)$, we obtain an Arrhenius-like representation:

$$\ln k = (c + \ln T) - \frac{\Delta H^\ddagger}{R} \cdot \frac{1}{T} \tag{6}$$

Consequently, the slope of the rate constant (plotted on a logarithmic scale) as a function of $1/T$ is approximately proportional to $\Delta H^\ddagger$. (The temperature dependence of the first term is small over a relevant temperature range and with the values of $\Delta H^\ddagger$ and $\Delta S^\ddagger$ assumed here.)
Figure S9 shows some exemplary cases.

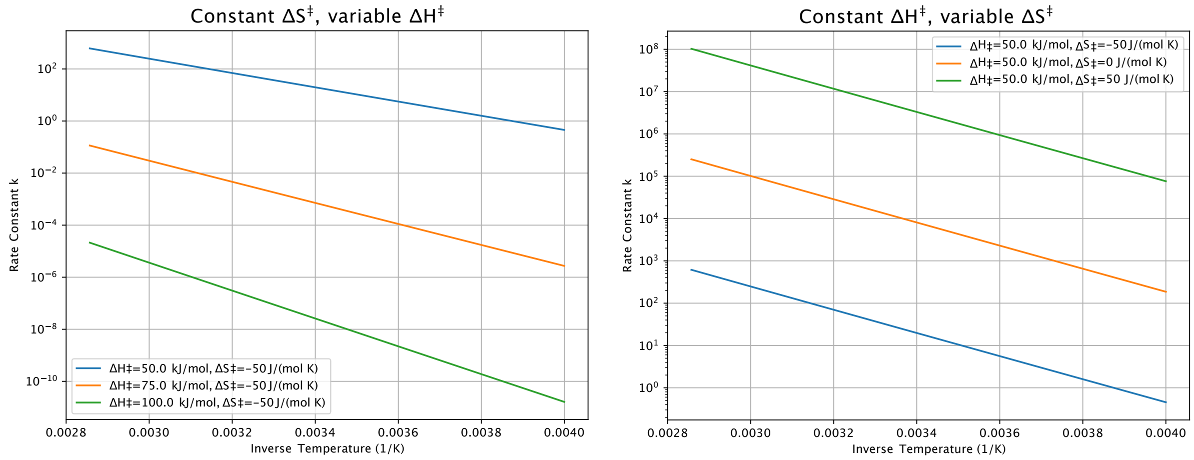

**Figure S9:** Model calculations of exchange rate constants with the Eyring equation, assuming different values of $\Delta H^\ddagger$ and $\Delta S^\ddagger$. This data illustrates that a flatter dependency of $\log k$ vs $1/T$ is due to smaller $\Delta H^\ddagger$. Moreover, these calculations support that the reduced rate constants in crystals is due to reduced $\Delta S^\ddagger$. The temperature range shown here is 250 to 350 K.

### 2 Supplementary Figures and Tables

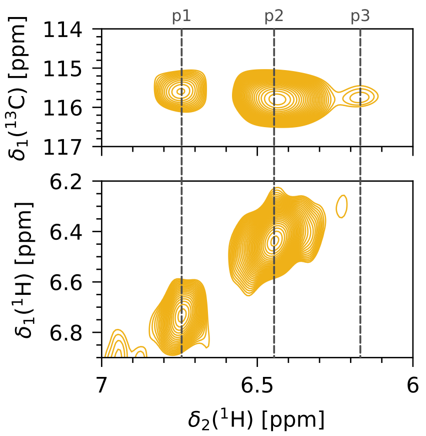

**Figure S10:** Dipolar-based $^{1}H-^{13}C$ 2D MAS NMR spectrum (upper panel) and $^{13}C-^{13}C$ EXSY spectrum with 256 ms exchange time and 2D $^{1}H-^{1}H$ readout (lower panel) of IgG:GB1$_{QDD}$. The $^{1}H-^{13}C$ spectrum is the same as the yellow spectrum in Figure 2D of the main text. The $^{1}H$ frequencies of p1, p2, and p3 are marked for better comparison to the EXSY spectrum. No cross-peaks can be observed in the EXSY spectrum, confirming that the three peaks belong to the three different tyrosines. If one of the tyrosines was in slow exchange and would therefore result in two signals, these two signals should show exchange cross-peaks in the EXSY experiment. For experimental details, see Table S2.

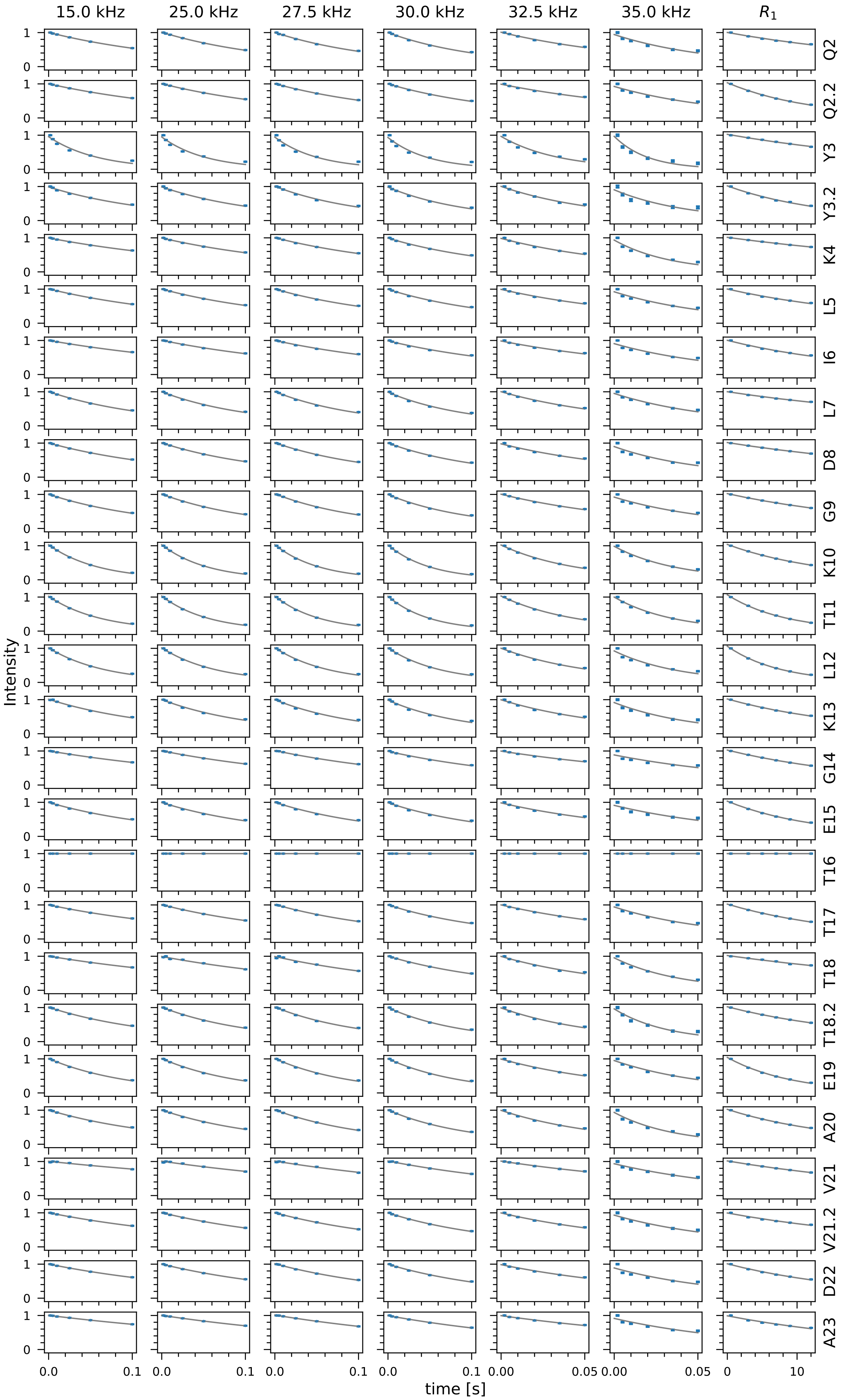

**Figure S11:** Continued on next page.

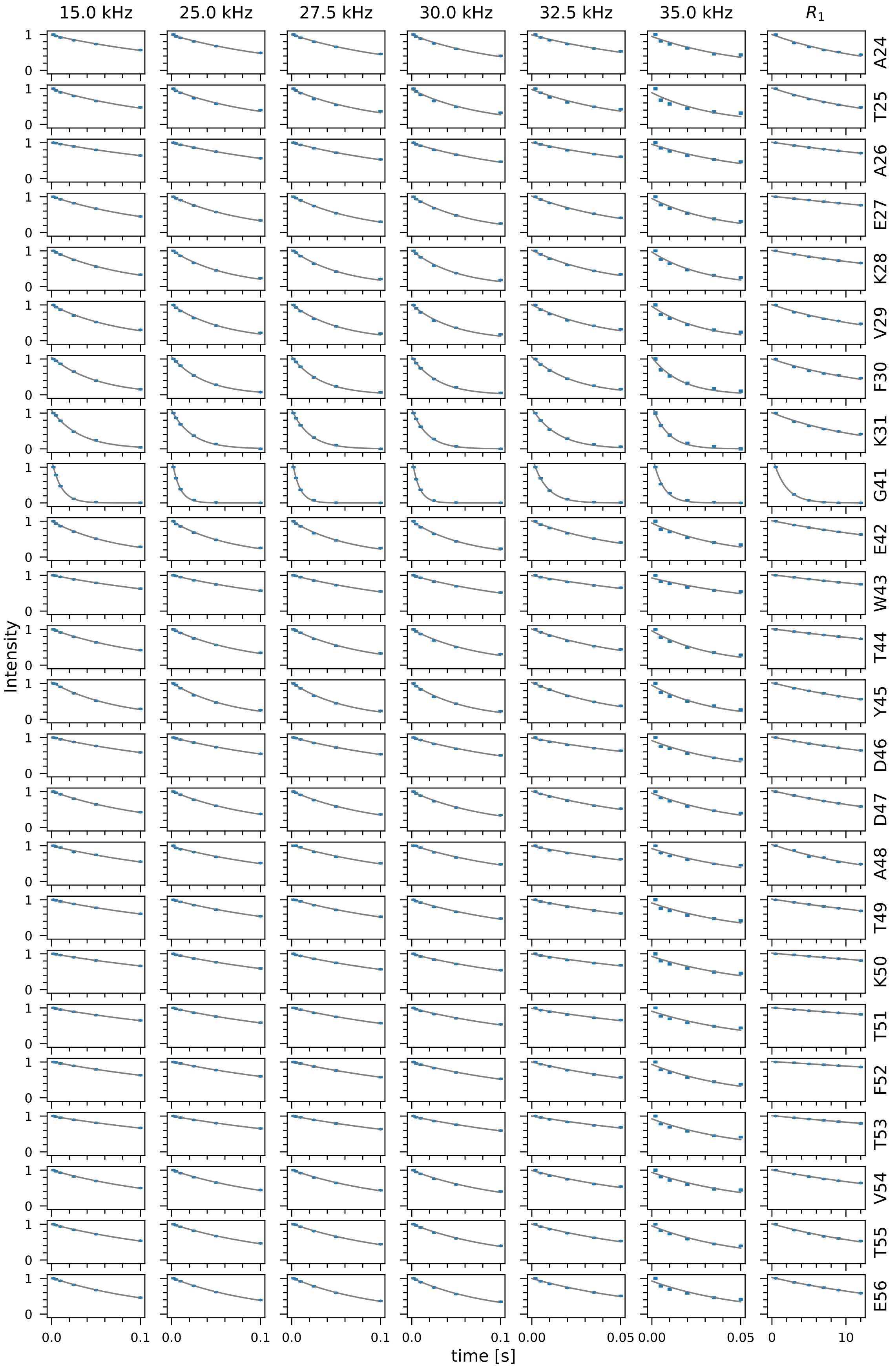

**Figure S11: Continued.** Exponential fits (gray) of experimental decay curves (blue) for $^{15}$N $R_1$ and $R_{1\rho}$ relaxation rate measurements of backbone amides in GB1$_{\mathrm{QDD}}$ crystals at a MAS frequency of 39 kHz and a temperature of 304 K. $R_{1\rho}$ was measured at $\nu_{\mathrm{SL}}$ = 15, 25, 27.5, 30, 32.5 and 35 kHz.

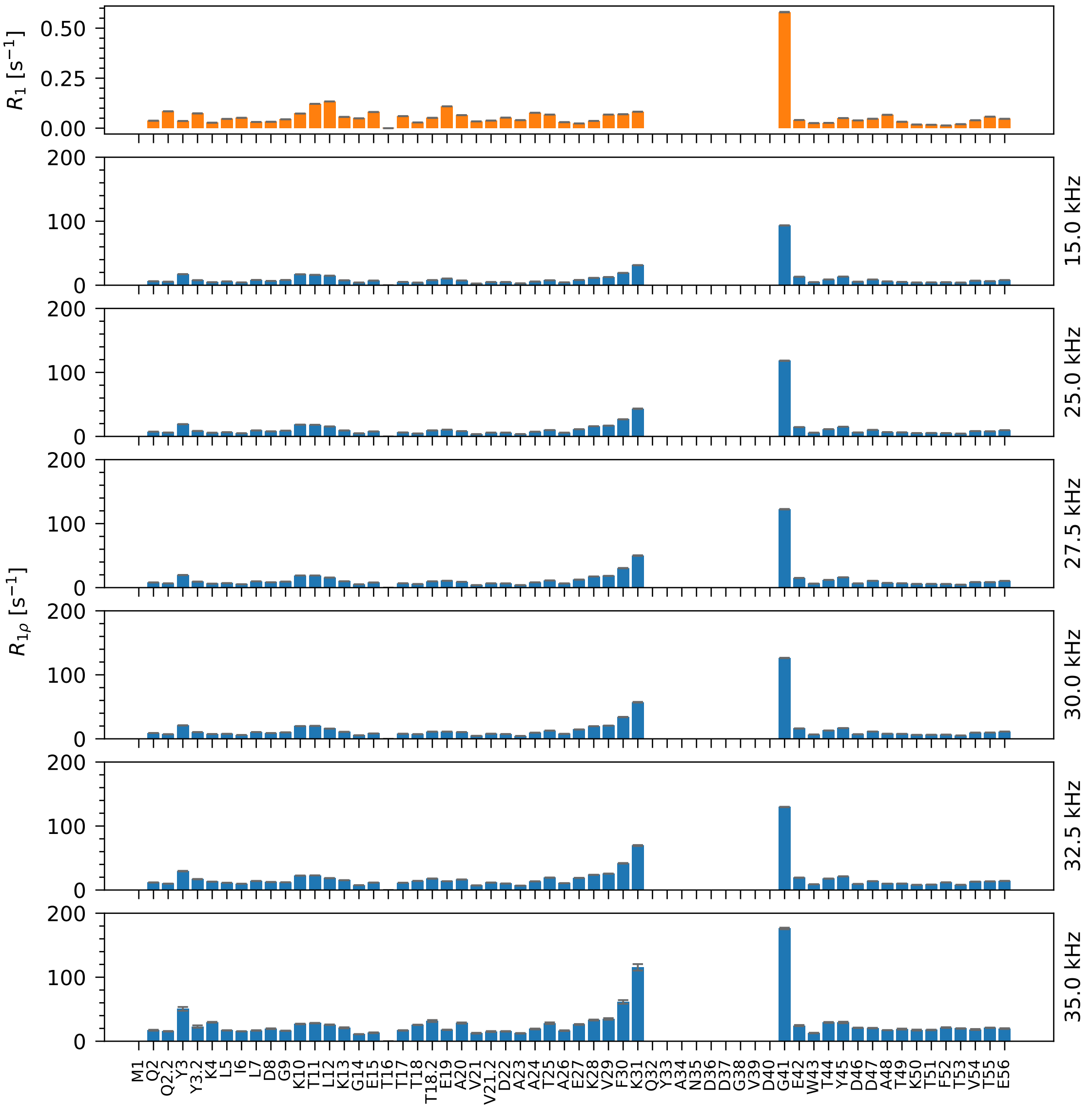

**Figure S12:** Residue-wise $^{15}$N $R_1$ (top panel) and $R_{1\rho}$ relaxation rate constants at $\nu_{SL}$ = 15, 25, 27.5, 30, 32.5 and 35 kHz as specified on the right side for crystalline GB1$_{QDD}$. Two values are reported for the residues that show two signals (Q2, Y3, T18, and V21).

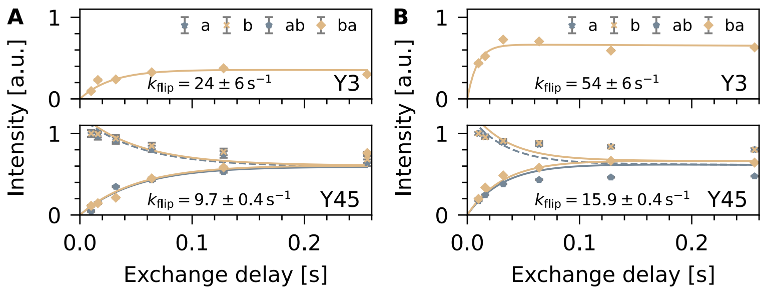

**Figure S13:** EXSY build-up and decay curves for Y3 and Y45. For Y3, only the build-up of 'ba' was fitted as the intensities of the other signals were strongly impacted by peak overlap with Y33. For Y45, we performed a combined analysis of all four peaks (*30*). (A) Measured at a MAS frequency of 39 kHz and a temperature of 288 K. (B) Measured at a MAS frequency of 30 kHz and a temperature of 304 K.

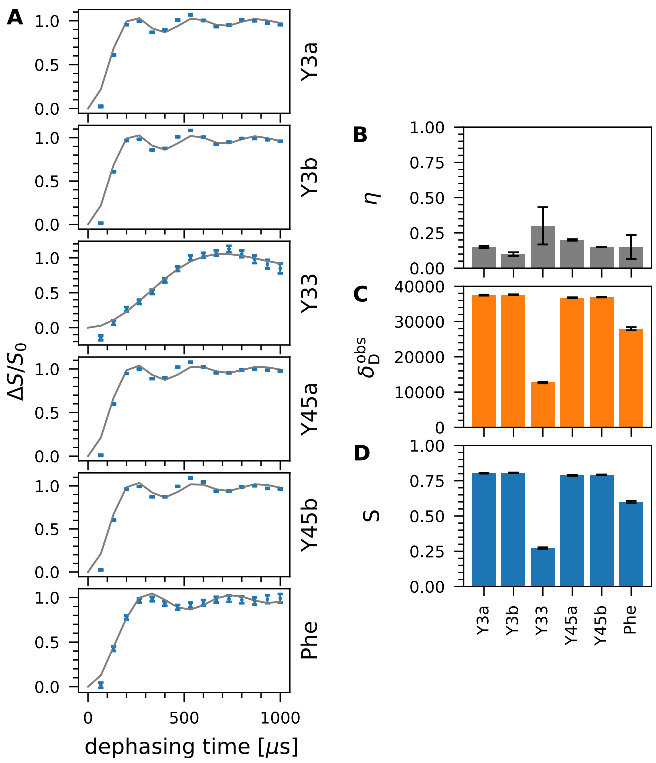

**Figure S14:** Determination of dipolar order parameters of $(CH)^{\epsilon}$-labeled tyrosines and phenylalanines in $GB1_{QDD}$ crystals. (A) Experimental (blue) and best-fit simulated (gray) REDOR curves. (B) Best-fit $\eta$. (C) Best-fit $\delta_D$. (D) Dipolar order parameter.

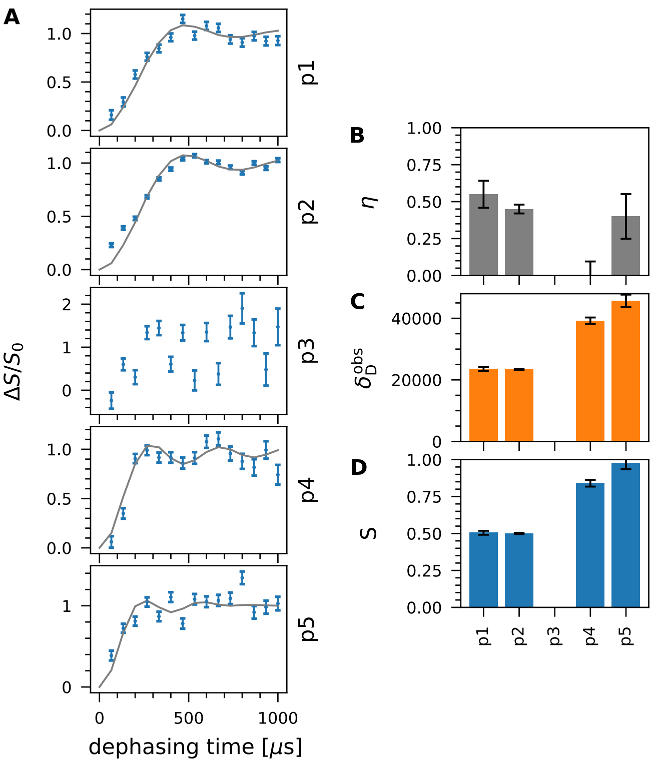

**Figure S15:** Determination of dipolar order parameters of $(CH)^{\epsilon}$-labeled tyrosines and phenylalanines in the IgG:GB1$_{QDD}$ complex. (A) Experimental (blue) and best-fit simulated (gray) REDOR curves. p3 has very low sensitivity which is why we did not fit the data. (B) Best-fit $\eta$. (C) Best-fit $\delta_D$. (D) Dipolar order parameter.

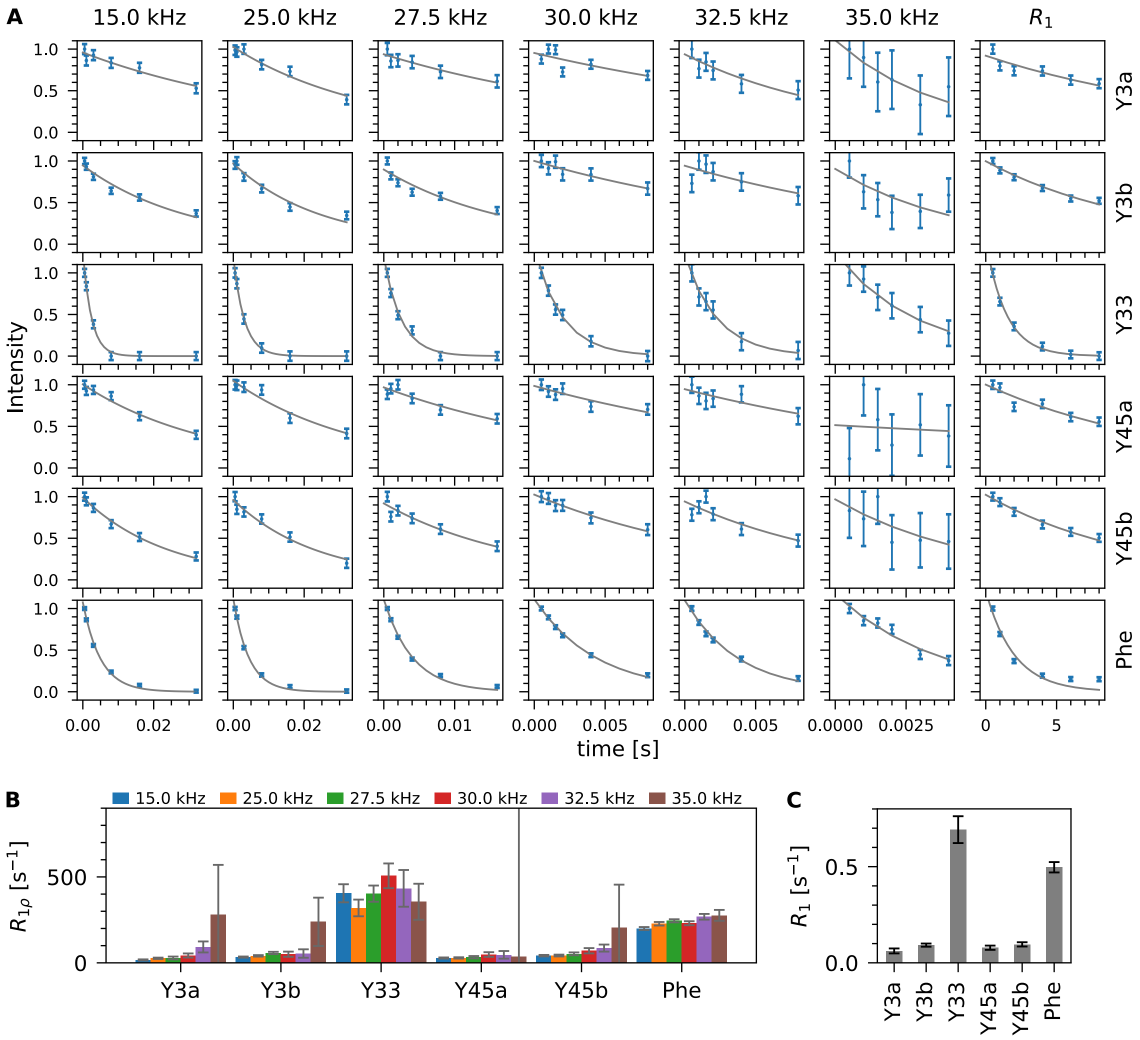

**Figure S16:** $^{13}$C $R_1$ and $R_{1\rho}$ relaxation rate constants of $(CH)^{\epsilon}$-labeled tyrosines and phenylalanines in GB1$_{QDD}$ crystals at a MAS frequency of 39 kHz and a temperature of 288 K. $R_{1\rho}$ was measured at $\nu_{SL}$ = 15, 25, 27.5, 30, 32.5 and 35 kHz. (A) Exponential fits (gray) of experimental decay curves (blue). (B) Fitted $R_{1\rho}$ relaxation rate constants at different spin-lock fields. (C) Fitted $R_1$ relaxation rate constants.

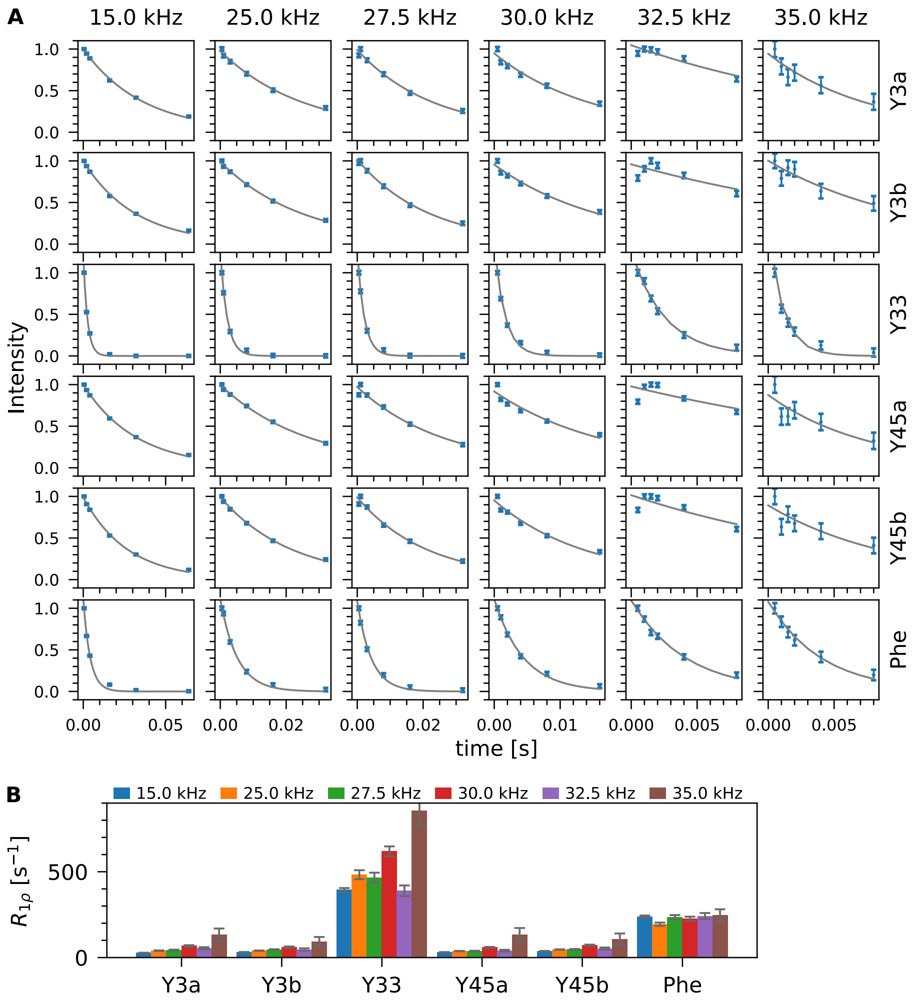

**Figure S17:** $^{13}$C $R_{1\rho}$ relaxation rate constants of $(CH)^{\epsilon}$-labeled tyrosines and phenylalanines in GB1$_{QDD}$ crystals at a MAS frequency of 39 kHz and a temperature of 304 K. $R_{1\rho}$ was measured at $\nu_{SL}$ = 15, 25, 27.5, 30, 32.5 and 35 kHz. (A) Exponential fits (gray) of experimental decay curves (blue). (B) Fitted $R_{1\rho}$ relaxation rate constants at different spin-lock fields.

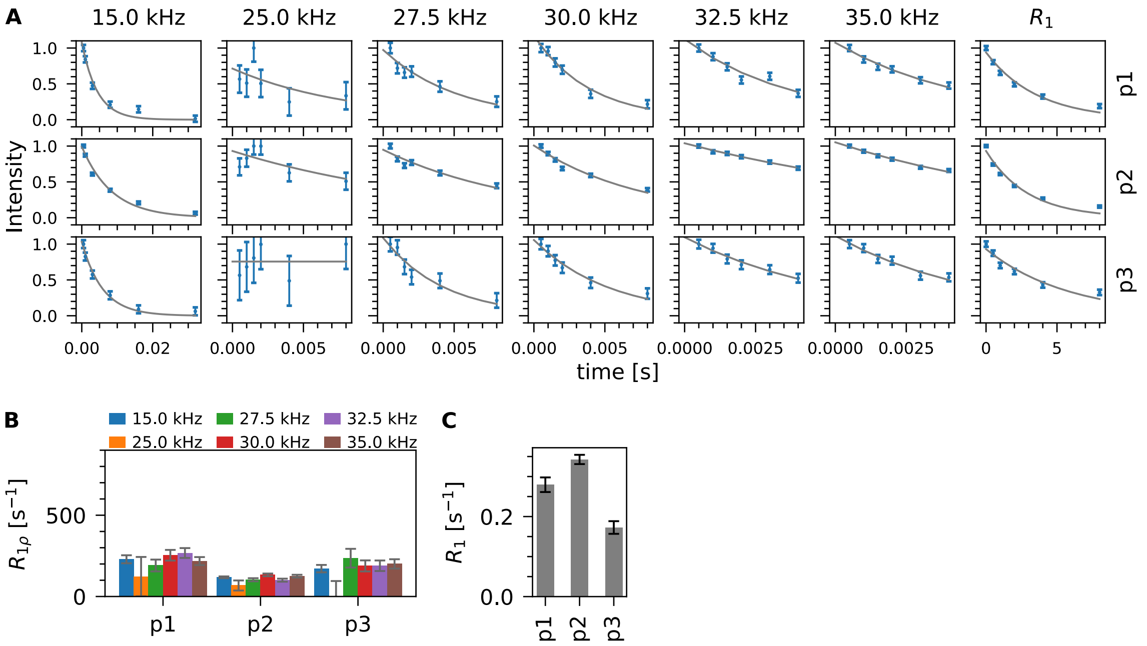

**Figure S18:** $^{13}$C $R_1$ and $R_{1\rho}$ relaxation rate constants of $(CH)^{\epsilon}$-labeled tyrosines and phenylalanines in the IgG:GB1$_{QDD}$ complex at a MAS frequency of 39 kHz and a temperature of 304 K. $R_{1\rho}$ was measured at $\nu_{SL}$ = 15, 25, 27.5, 30, 32.5 and 35 kHz. (A) Exponential fits (gray) of experimental decay curves (blue). (B) Fitted $R_{1\rho}$ relaxation rate constants at different spin-lock fields. (C) Fitted $R_1$ relaxation rate constants.

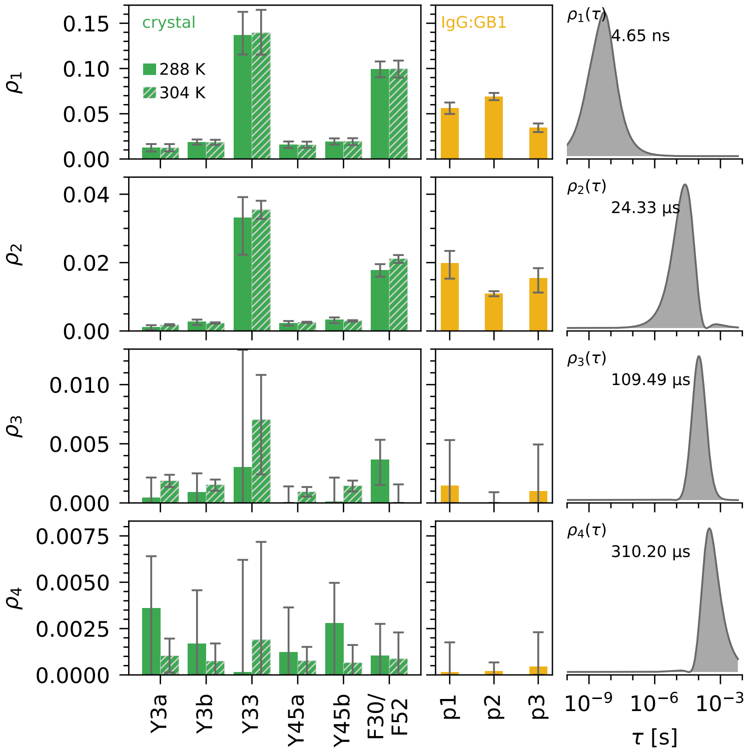

**Figure S19:** Detectors analysis for $(CH)^{\epsilon}$-labeled tyrosines and phenylalanines in $GB1_{QDD}$ crystals (green) and in the IgG:$GB1_{QDD}$ complex (yellow). The detector sensitivities $\rho_i(\tau)$ are shown on the left. The correlation time corresponding to the maximum of $\rho_i(\tau)$ is specified. The bar plots show the corresponding responses $\rho_i$ for each residue for the two samples. The crystal was measured and analyzed at two temperatures. $R_1$ was only measured at the lower temperature but included in both analyses as it is not expected to be very temperature-dependent in our available temperature range.

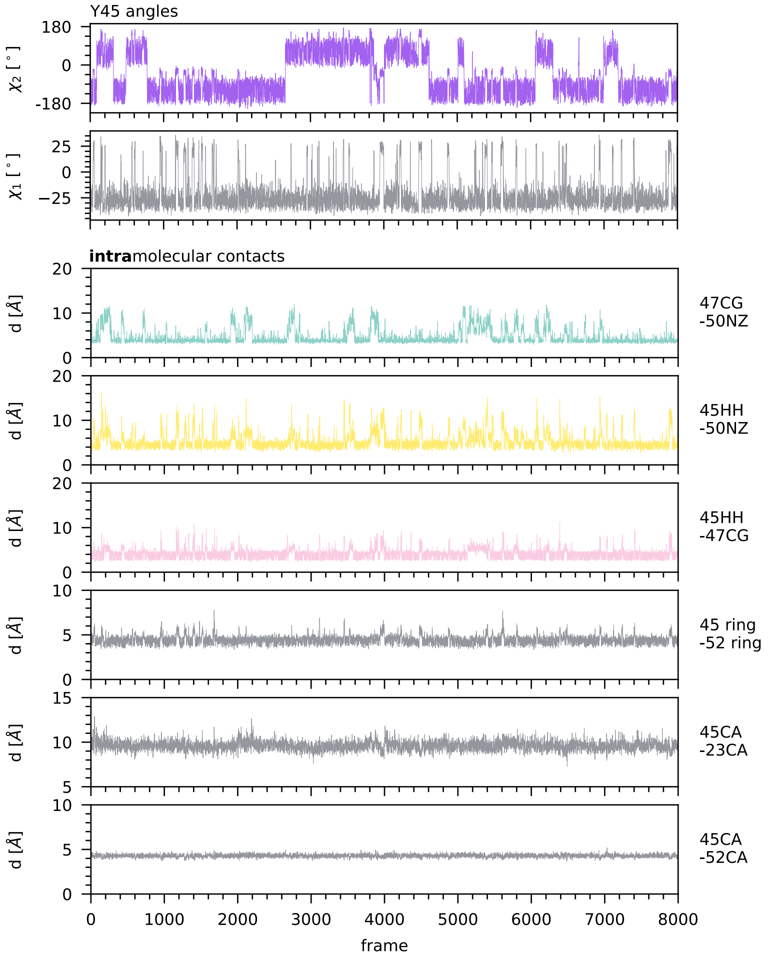

**Figure S20:** Traces of selected angles and distances from solution MD simulations. Distances that are not visualized in Figure 6A-B of the main text are shown in Figure S22.

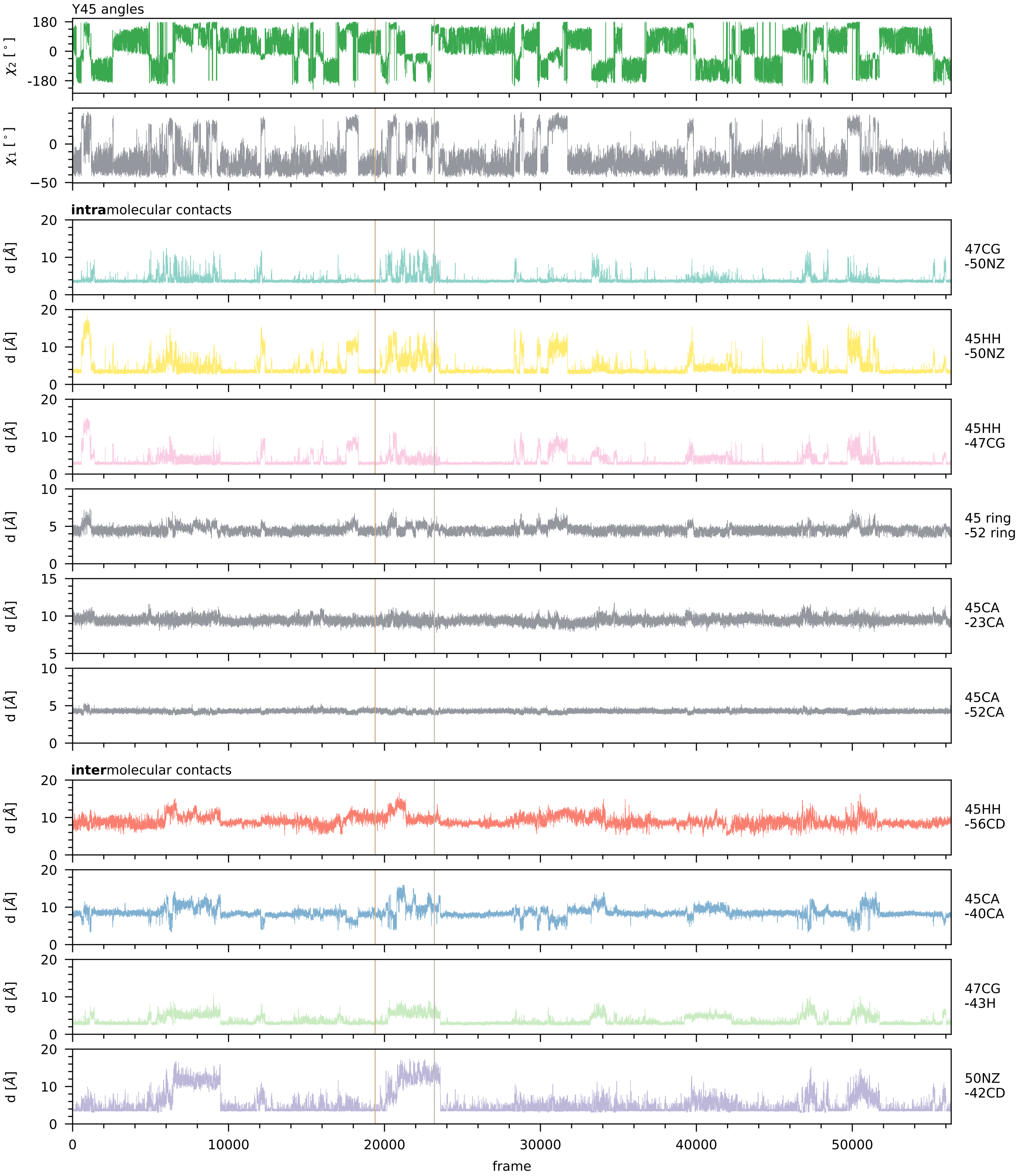

**Figure S21:** Traces of selected angles and distances from crystal MD simulations. Sections of these traces are shown in Figure 6G of the main text. Distances that are not visualized in Figure 6A-F of the main text are shown in Figure S22.

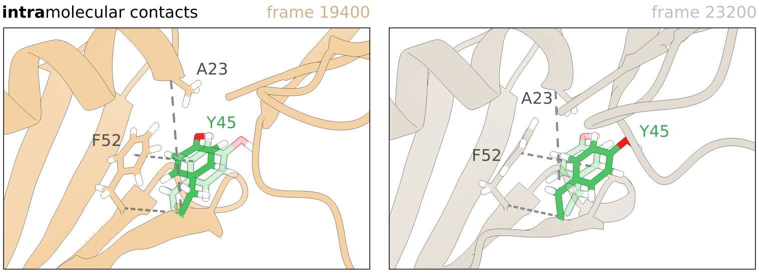

**Figure S22:** Visualization of distances whose traces are shown in Figures S20 and S21 but not in Figure 6 of the main text.
